# Bias invariant RNA-seq metadata annotation

**DOI:** 10.1101/2020.11.26.399568

**Authors:** Hannes Wartmann, Sven Heins, Karin Kloiber, Stefan Bonn

## Abstract

Recent technological advances have resulted in an unprecedented increase in publicly available biomedical data, yet the reuse of the data is often precluded by experimental bias and a lack of annotation depth and consistency. Here we investigate RNA-seq metadata prediction based on gene expression values. We present a deep-learning based domain adaptation algorithm for the automatic annotation of RNA-seq metadata. We show how our algorithm outperforms existing approaches as well as traditional deep learning methods for the prediction of tissue, sample source, and patient sex information across several large data repositories. By using a model architecture similar to siamese networks the algorithm is able to learn biases from datasets with few samples. Our domain adaptation approach achieves metadata annotation accuracies up to 12.3% better than a previously published method. Lastly, we provide a list of more than 10,000 novel tissue and sex label annotations for 8,495 unique SRA samples.

## Introduction

Next generation RNA-sequencing (RNA-seq) has been a pillar of biomedical research for many years (1, 2). It allows researchers to simultaneously quantify and compare the expression of tens of thousands of genomic transcripts. A continuous drop in cost makes RNA-seq a widely available method of choice to uncover the molecular basis of biological development or disease (3, 4). As a result of this, recent years have seen a strong growth in publicly accessible RNA-seq data. The actual reuse and integration of this data, however, has been largely limited by the lack of consistent metadata annotation and individual dataset bias (5, 6). The lack of metadata annotation for RNA-seq samples, such as tissue of origin, disease or sex phenotype, prohibits experimenters from finding data that is relevant to their research. Moreover, dataset biases (7) due to differences in protocols and technologies (8) or of a biological nature hinder integration and comparative analysis.

To allow for efficient data reuse, publicly available data has to be harmonized and well annotated with standardized meta-data and subsequently be made accessible (and searchable) (9); This practice is followed by the Genotype-Tissue Expression Project (GTEx) (10), and The Cancer Genome Atlas (TCGA). Nevertheless, the primary database for next-generation sequencing projects, the Sequence Read Archive (SRA) (11), stores raw sequencing information that lacks rigorous standards of curation, which limits the reusability of its data.

Efforts to predict missing or sparse metadata in public RNA-seq resources have shown promising results. For instance, recently published studies used text mining approaches to retrieve missing annotation from associated abstracts or free text annotations in the data sources (12–14). Others have used RNA-seq expression values for phenotype prediction. For example, machine learning (ML) has successfully been applied to disease and cell type classification (15, 16) or survival outcomes on TCGA data (17). Others have taken advantage of prior domain knowledge such as gene regulatory networks for enhanced feature selection (18, 19). Recently a linear regression model fitted to GTEx data has been presented for the prediction of tissue, sex and other phenotypes of SRA and TCGA samples (20). These efforts provide evidence that missing RNA-seq metadata can be successfully predicted based on genomic expression values using ML approaches.

Artificial neural networks (ANNs) in their various forms and functions consistently outperform classical ML approaches in a large variety of biological tasks, including classification, data generation and segmentation (21–24). Given large training datasets these algorithms can learn complex representations of data by automatically weighting and combining features non-linearly. This has led us to hypothesize that ANN based models could increase the performance in meta-data prediction beyond that of classical ML approaches such as linear regression. Of special interest in this context is domain adaptation (DA) (25), a subfield of ML which aims to specifically alleviate problems conferred by dataset bias (26). The aim of DA is to build and train ANNs on a source domain in such a way that the model performs well on a biased target domain. One strategy that has been successfully applied is interpolation between source and target domain by training feature extractors on an increasing ratio of target to source domain data (27). Another popular strategy is adversarial training by applying two loss functions. The first loss function forces the model to learn weights for class prediction while the second forces the model to learn to ignore differences between the source and target domain (28). Tzeng et al. adapted this idea (29), proposing a model using a separate source and target encoder using them as ‘real’ and generator input for a generative adversarial network (30) that is capable of ignoring bias. For the case of scarce target data a similar approach was previously proposed using siamese networks (31, 32). All these methods have been implemented and applied by us for RNA-seq phenotype prediction and found not to be scalable to a situation with hundreds of different and scarce target domains, encountered, for instance, in the SRA. Here we present a DA approach capable of leveraging a number of dataset biases, boosting generalizability of phenotype prediction. We developed the model using three data sources (GTEx, TCGA and SRA) of different size and with a different degree of bias. To validate our approach we compare it to a previously suggested linear model (LIN) (20) as well as a standard supervised multi-layer perceptron (MLP) on prediction of tissue of origin, sex and sample source. Importantly, we find that our DA network significantly outperforms the strongly supervised LIN model by up to 12.3% in prediction accuracy. We subsequently apply trained models to generate and make available new metadata for 8,495 unique SRA samples.

## MATERIALS AND METHODS

### Data Acquisition

To train and test models we gathered data from three different sources, each with a different level of homogeneity, which we define as the number of unique dataset biases present within one data source (Supplementary Figure 1). Biases stem from the unique circumstances, protocols and reagents used as well as biological factors of the study (7, 8). Here we define a dataset as all the RNA-seq samples from one study based on the assumption that they were obtained and processed under equal conditions. To avoid additional biases by the use of different bioinformatic alignment pipelines (35) all data was downloaded from recount2 (release 13.09.19, https://jhubiostatistics.shinyapps.io/recount/). Recount2 aggregates raw RNA-seq data from different sources and re-runs the data through the Rail-RNA alignment pipeline (36). The RSE V2 files of all available RNA-seq projects (n=2,036) from recount2 were downloaded using the recount R package (v 1.11.13). The downloaded data was separated into three different data sources according to their origin. Figure 1A gives a general overview of the data obtained, the pre-processing steps and data set preparation.

**Fig. 1.**
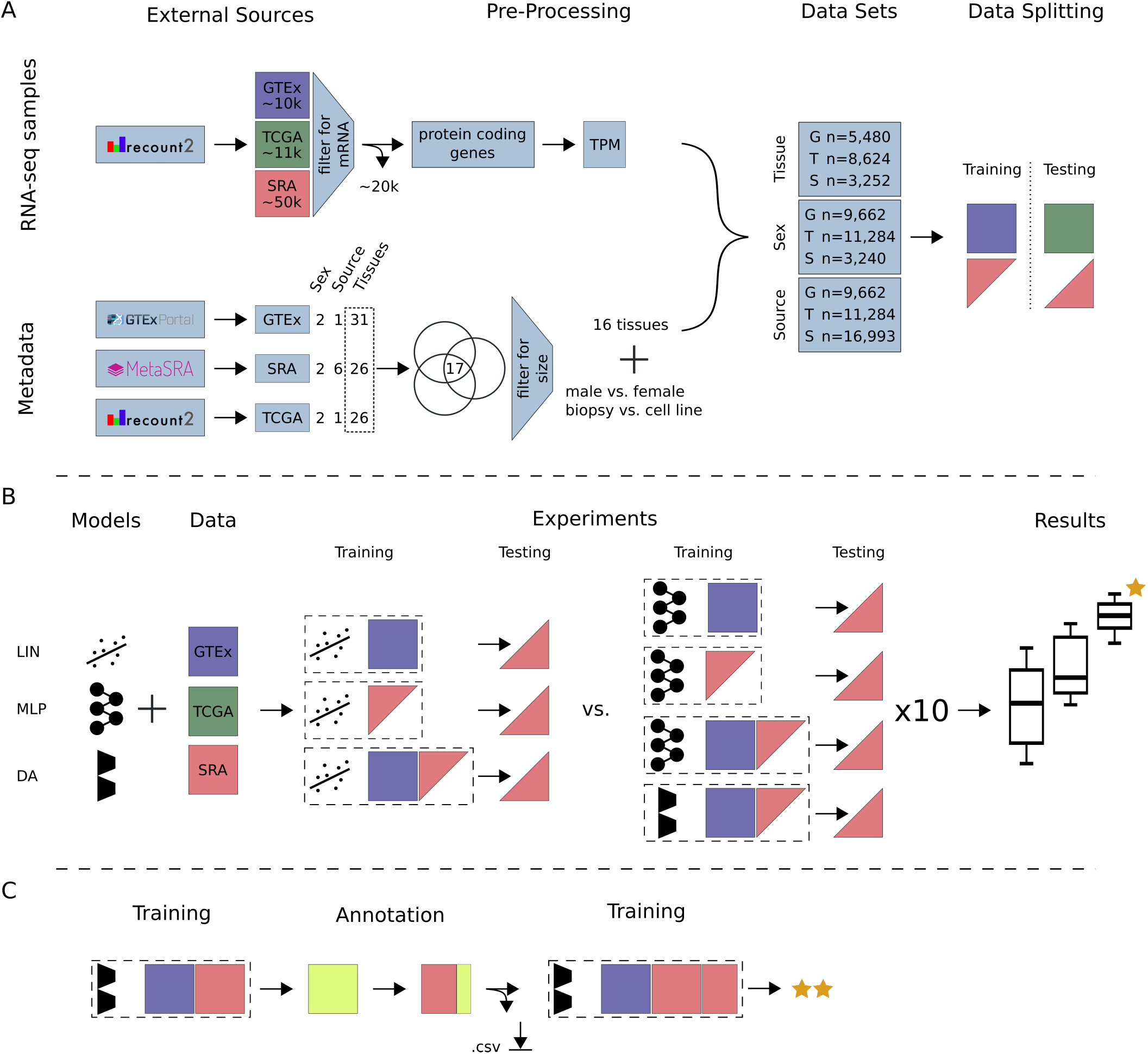
Study Overview. (A) All data available on recount2 was downloaded and split into three data sources: (i) GTEx, (ii) TCGA and (iii) SRA. Single-cell and small RNA samples as well as technical replicates were removed from the SRA data. Protein coding genes were selected from the gene count tables and TPM normalized. Metadata for tissue of origin (e.g. heart), source (e.g. biopsy) and sex phenotype was collected, if available. A subset of 17 tissues (common to GTEx, TCGA and SRA) was selected and filtered for class size, resulting in 16 tissue classes. For sample source the two largest classes in SRA were selected. Samples were subsequently annotated and training and testing data sets were created. GTEx was only used for model training unless stated otherwise. TCGA was only used for model testing. SRA was split such that samples from one study are exclusively in the train or test set. (B) We compare three models: LIN (linear model), MLP (multi-layer perceptron) and DA (novel domain adaptation algorithm). Experiments are different combinations of models and data sources. Here, an exhaustive list of experiments for tissue and sex classification tested on SRA data is depicted. Each configuration (dashed box) is made up of a model and training data. The previously published LIN model served as a benchmark for our MLP and DA model. Each model configuration was trained 10 times with different seeds to give an estimation of uncertainty. The best model (orange star) was chosen by comparing average performance across all seeds. (C) After determination of the best model, all available data was used for model training. Previously unlabeled SRA data (yellow square) was automatically annotated with the appropriate metadata. Newly annotated metadata can be used to re-train existing models to further improve performance. A list of all new metadata can be downloaded with the Supplementary Material.

### GTEx

The Genotype-Tissue Expression Project v6 (https://www.gtexportal.org/) comprises 9,662 samples from 554 healthy donors across 31 tissues. GTEx strives to build a highly homogeneous dataset with strict guidelines on donor selection, biopsy and sequencing methodology (more information at: https://www.gtexportal.org/home/documentationPage). We considered the GTEx data source to have a single dataset bias.

### SRA

From the Sequencing Read Archive, a total of 2,034 studies containing a total of 49,657 samples were downloaded from recount2. Every SRA study was potentially processed at a different site by a different technician following different standards. In addition, the underlying biological condition of the samples is often unclear. We assume each study to have a unique dataset bias which makes the SRA a highly heterogeneous data source. In addition, data annotation is not standardized resulting in sparse metadata with low fidelity.

### TCGA

RNA-seq data for The Cancer Genome Atlas (https://www.cancer.gov/about-nci/organization/ccg/research/structural-genomics/tcga) was downloaded consisting of 11,284 samples spanning 26 tissues. While there are 740 samples of healthy donors across 20 tissues, more than 90% of the samples are tumor biopsies from different tissues and different stages of tumor progression. TCGA accepts sequence data from different locations using different sequencing technologies. Despite the high level of standardization and reliability of metadata information, heterogeneity is also inherent to the TCGA dataset due to the biological context (cancers, stages) albeit not as pronounced as in SRA.

### Preprocessing of SRA Data Source

In this study we focus on bulk mRNA-seq data, as it is by far the most frequent RNA type in either of the three data sources used. The following approaches were used to remove data from single-cell and small RNA-seq studies from further analysis: First, we identified small RNA-seq data on the basis of the total fraction of small RNA counts and protein coding RNAs. Specifically, we considered a subset of the Gencode gene types (i.e. protein_coding and processed_pseudogene vs. rRNA, miRNA, misc_RNA, snRNA and lincRNA). Every sample that had its maximum total count fraction not allocated to either protein_coding or processed_pseudogene was removed from further analysis (Supplementary Figure 2). Second, we removed single-cell RNA-seq studies by scanning titles and abstracts for variations of the words ‘single cell’ and manually validated and excluded the identified samples. In addition to this semi-automatic validation step we manually validated the 50 largest projects within the SRA data source and removed samples that did not qualify as bulk RNA-seq data. Most importantly, we noticed a large number of technical replicates in the remaining SRA data. Using technical replicates to train and test a classification model inflates the reported metrics. Therefore only samples with a unique experiment accession (SRX) were retained. From the 49,657 SRA samples downloaded initially, 29,685 samples and 1,833 unique studies passed our preprocessing steps.

### Metadata

We considered three different phenotypes for expression based prediction. Explicitly, we predicted the tissue of origin of a biopsy (e.g. heart, lung, kidney, ovary), the patients’ sex, and sample source (denoting whether the sample was from a patient biopsy or a lab grown cell line) (Figure 1A).

### GTEx and TCGA

Tissue and sex annotation for GTEx were extracted from the official sample annotation table as provided by GTEx (GTEx_Data_V6_Annotations_SampleAttributesDS.txt, from https://storage.googleapis.com/gtex_analysis_v6/annotations). An annotation file for TCGA was provided by recount2. For tissue and sex annotation we took columns gdc_cases.project.primary_site and gdc_cases.demographic.gender respectively. Sample source was assumed to be of type biopsy for all GTEx (n=9,662) and TCGA (n=11,284) samples.

### SRA

For the SRA samples we relied on normalized meta-data provided by MetaSRA (14). Available SRA identifiers were downloaded through the GUI on http://metasra.biostat.wisc.edu by searching for all 31 GTEx tissues (site accessed on 11.09.2019). Supplementary Table 1 lists assumed mappings from GTEx tissue names to MetaSRA tissue names where no direct mapping was available. Of the 31 tissues available for GTEx we were able to identify samples for 26 in MetaSRA, resulting in 6,183 annotated SRA samples. Sample identifiers for sex were accessed through the same GUI by searching for male organism and female organism + Homo sapiens cell line which resulted in 3,240 annotated SRA samples. Sample source was determined using the sqlite file provided by MetaSRA (metasra.v1-5.sqlite, http://metasra.biostat.wisc.edu/download.html, colum sample_type) resulting in 28,043 annotated samples across six sample source categories.

### Tissue Label Harmonization

GTEX, TCGA and SRA have 17 common tissue types (Supplementary Figure 3). Bladder was removed due to its small sample size (GTEX n=11). We kept samples of comparable size in SRA (adrenal gland n=14, testis n=14, pancreas n=17 in the SRA training data), as the SRA training data is mainly used for bias injection, such that size was not considered an exclusion criterion. This resulted in 5,480, 8,624, and 3,252 tissue annotated samples across 16 tissues for GTEx, TCGA and SRA, respectively (Supplementary Tables 2 and 3).

### Dimensionality Reduction and Normalization

The downloaded gene count table provided counts for 58,037 genes (Gencode v25, GRCh38, 07.2016). First standard log2 Transcript per Million (TPM) normalization was applied to normalize for gene length and library size. We next reduced the number of input features (genes), aiming to keep features that contain information and removing potentially uninformative features. First, all non-protein coding genes were removed, reducing the gene set by 65.5% to 19,950 genes. For sex classification only protein coding genes on the X and Y chromosome (n=913) were selected. For retaining only genes that show significant dispersion across tissues, we computed the Gini coefficient (15, 37, 38) for all remaining genes across all GTEx samples. Housekeeping genes, for example, are known to be expressed similarly across tissues and would score a low Gini coefficient (i.e. high dispersion). Low and high cutoffs were applied during hyperparameter optimization. For tissue classification, genes with Gini coefficients *g* between 0.5 and 1 were retained, resulting in a features space of dimension d=6,974. For sex classification, genes with 0.4 *< g <* 0.7 were used (d=190). Sample source classification included genes with 0.3 *< g <* 0.8 (d=8,679) (Supplementary Table 2).

### Dataset Preparation

#### Phenotype Classification Experiments

**Tissue**: To ensure that dataset biases are not shared between train and test sets, SRA data was always split on the study level. For tissue of origin prediction the two largest SRA studies per class were put in the training set. This ensured maximal bias variability in the remaining test data, ensuring a realistic test score. Of the 178 SRA studies containing tissue annotated samples, 30 studies were selected for the training set (n=1,721) and 148 studies for the test set (n=1,531) (Supplementary Tables 2 and 3). **Sex**: In total, 159 SRA studies contained samples annotated with male and or female by MetaSRA. These studies were combined into the training set (studies=78, n=2,317), and test set (studies=81, n=923) (Supplementary Tables 2 and 3). For model validation GTEx was randomly split into training and test sets with a 80:20 ratio for both sex and tissue classification. **Sample Source**: A confidence cutoff of ≥ 0.7 was applied (provided by MetaSRA), reducing the total amount of annotated samples for SRA from 23,651 to 17,343. For each of the two selected SRA categories (i.e. biopsy and cell line) we sorted all available studies by number of samples, placed the first third of studies into the training (studies=420, n=12,725), the second third into the test (studies=422, n=3,144) and the last third into the SRA validation set (studies=418, n=1,124) (Supplementary Tables 2 and 3). A list of the sample ids and corresponding labels is available in the Supplementary Material.

#### Metadata Annotation

After determining the best model for each phenotype we re-trained the models for automated metadata annotation. The same datasets as defined above were used for the sex metadata annotation. **Tissue**: We followed the same pipeline as described above, the only difference being that no samples were discharged because of their tissue label. Samples from a tissue class other than the original 16 classes were pooled together into a ‘catch-all’ class, resulting in 17 classes. In total 44 SRA studies were selected for the training set (n=3,370) and 203 studies for the test set (n=2,813). **Sample Source**: Contrary to before, for metadata annotation we used all available classes in the SRA data source. All classes that are not tissue (i.e. biopsy) were grouped into a single ‘catch-all’ class while the same cutoff as before was applied. The training set (n=16,463) is made up of 974 SRA studies and the test set (n=3,707) of 492 studies.

#### Multilayer Perceptron

**MLP.** MLPs use fully connected neural network layers to learn non-linear features from a raw input space (33) and constitute the most basic form of ANNs. All our ANN based models were developed and trained on tf.keras (Tensorflow 2.1). The hyperparameters for each prediction task were determined using exhaustive iterative random search (keras tuner 1.0.1) (Supplementary Table 4). In case of approximately equal accuracy on the validation set, the least complex model was chosen. A single hidden layer was used in each case with 128, 128 and 32 nodes for tissue, sample source, and sex prediction, respectively (Supplementary Table 5, Supplementary Figure 4). Each network was trained for 10 epochs with a batch size of 64. Performance was quantified by mean sample accuracy and mean class accuracy and subsequently used to benchmark our DA approach.

#### Domain Adaptation Model

**DA.** Many DA models correct bias between two domains, a source and a target domain. In biological research, however, one is often confronted with a large number of small datasets, each potentially with its unique dataset bias. Therefore, we specifically designed our DA model to be able to learn from very few data by using a siamese network architecture (31). The siamese network learns bias from pairs or triplets of training samples by exposing each sample in multiple relationships to the model. We distinguished three different types of input data for our model. The source domain is a large single-bias dataset used to learn the feature embedding for the classification task (in our case: GTEx). The bias domain contains labeled samples from multiple smaller datasets (in our case: SRA) each with its own bias. The target domain refers to unlabeled and biased datasets we want to classify (unlabeled SRA or TCGA data).

#### Model Architecture

Our DA architecture is based on the siamese network architecture. It consists of three modules: A source mapper (SM) and bias mapper (BM) which correspond to the siamese part of the model, as well as a classification layer (CL). These modules give rise to three different configurations, i.e. two training cycles and a prediction configuration (see Supplementary Figure 5 for a brief illustration). In the first training cycle the source mapper (SM) and the classification layer (CL) are combined to form an MLP (Figure 2A). The task of the SM is to learn a mapping from the input space to an embedding space from which the CL can predict phenotype classes. The SM-CL module is trained with a batch size of 64 for 10 epochs. Because the SM-CL MLP is trained on a large single-bias dataset it will likely overfit and thus not readily generalize to other datasets (Figure 2B). For a second training cycle, the bias mapper is created with the same architecture as the SM. The CL is removed and the weights of the SM are frozen. Triplets of data are forward propagated through the BM and SM in parallel (Figure 2C). Each triplet is made up of an anchor (a) sampled from the bias domain, and a positive (p) and a negative sample (n) from the source domain. The anchor and the positive sample have equal class labels whereas the negative sample is from a randomly selected different class. The triplets loss function (34) was used to optimize the model during training:

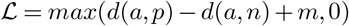

Where *d*(*i, j*) are the distances in embedding space between the respective outputs of the BM and SM on samples i and j. For improved training time and robustness, our model is trained on semi-hard triplets (34)

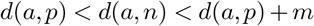

with a margin parameter m. The distances are defined as Euclidean distances in embedding space:

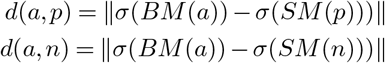

*σ* is the sigmoid activation function for the embedding vector. The SM-BM module was trained for 10 epochs with a batch size of 64. Hyperparameters were determined as described above (Supplementary Table 5, Supplementary Figure 4). As this training cycle proceeds, the BM learns to map its output onto the SM embedding space. After training, the bias mapper and the classification layer are combined to a BM-CL MLP and can be used for prediction of the target domain (Figure 2D). The source code as well as an example are available at: github.com/imsb-uke/rna_augment.

**Fig. 2.**
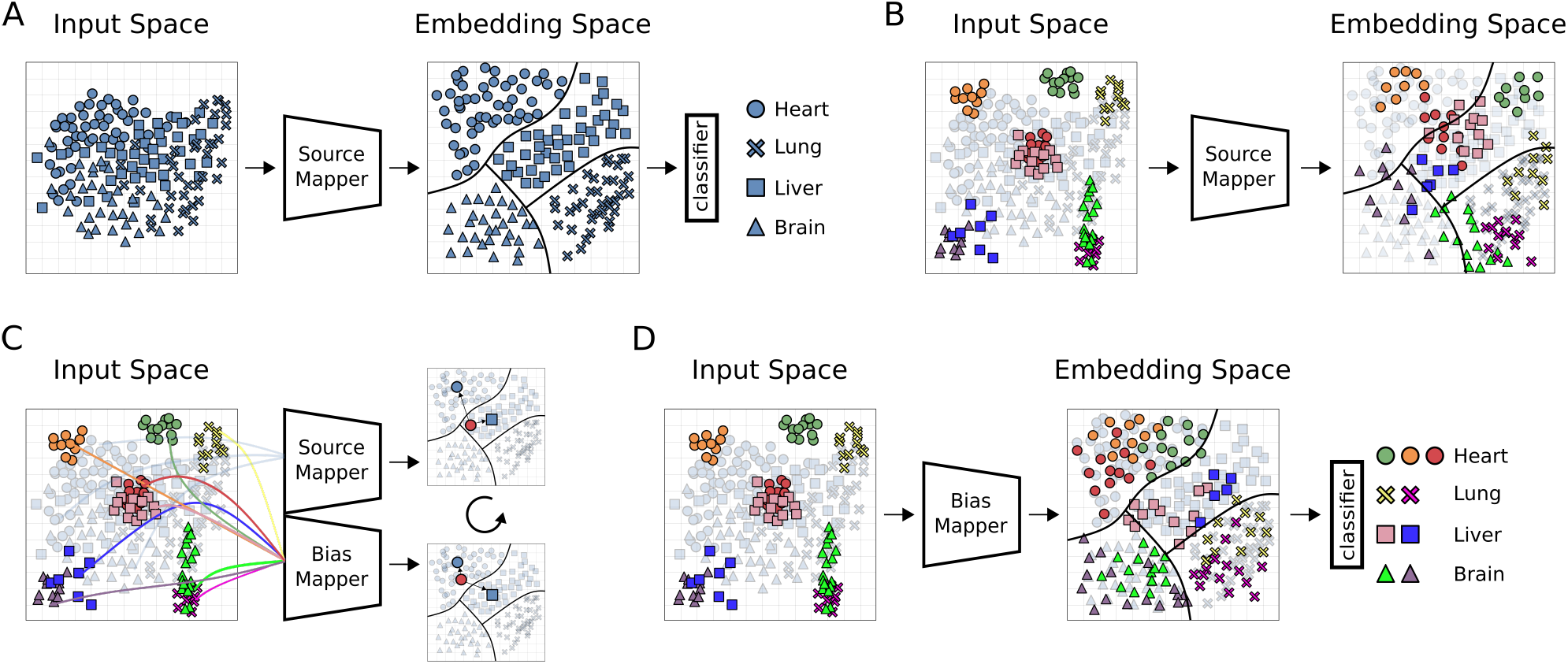
Overview Domain Adaptation Model. Illustration of our DA model architecture and training. Shapes of (hypothetical) data points represent classes, colors are datasets with unique biases. Source Mapper (SM), Bias Mapper (BM) and classifier layer (CL) are ANN modules. (A) First training cycle: The SM is trained on a single bias dataset, the source domain (SD). In this step, the SM learns a feature embedding. The CL learns how to partition this embedding space into classifiable regions and draws decision boundaries (black lines). (B) For biased test data (colored sample data points), same classes may occupy distinct regions in input space. In this case, the source mapper may not be able to map the samples to the correct region of embedding space, compromising classification performance of the CL. (C) In order to learn the mapping of different biases to the embedding learned in (A), a bias mapper (BM) is created by copying the SM, and trained weights of the SM are fixed. In this second training cycle, triplets of samples are passed through the SM-BM configuration, consisting of an anchor from the bias domain and two samples from the source domain, one of them with a matching label. The triplet loss function is defined to minimize distance of like labels in embedding space and to maximize distance of opposite labels. This process is repeated until the SM has learned to map all known biases into the previously learned embedding space. (D) The BM is now able to map data points from previously unseen datasets into the embedding space where the CL can classify them.

#### Linear Regression Model

**LIN.** We used the metadata prediction performance of the LIN model described in Ellis et al. (20) as point of reference. The LIN model was optimized on the same data as all other models (see data section of methods). For each experimental setup, the following steps were conducted in R version 3.6.3 in order to build the corresponding phenotype predictor and evaluate its accuracy based on the test data:

1. calculating the coverage matrix for the training samples based on the regions reported in Ellis et al. (20) by employing the function ‘coverage_matrix_bwtool’ (R package recount.bwtool version 0.99.31).
2. building the model by running ‘filter_regions’ and ‘build_predictor’ (R package phenopredict version 0.99.0) with the same parameters used in Ellis et al. (20)
3. testing the model on the test samples with ‘extract_data’, ‘predict_pheno’, ‘test_predictor’ (R package phenopredict version 0.99.0)

Notably, our experiments differ from the original work (20) solely by applying additional preprocessing steps to the samples (see Methods), which may be responsible for observed small differences in performance. For implementation details and code examples for the before-mentioned functions, see the documentation (http://rdrr.io/github/ShanEllis/phenopredict/).

#### Nomenclature of Experiments

Each experiment was named after the model, the training and the test data used. The possible models are LIN (linear model (20)), MLP (multi-layer perceptron) and DA (novel domain adaptation approach). The data sources are named G (GTEx), T (TCGA) and S (SRA). If only the SRA training data is used (i.e. if the model is evaluated on the SRA test data) we write S_small_. If the SRA train and test sets are combined for training we write S_large_. For instance, an experiment using an MLP, trained on a mix of GTEx and SRA and evaluating on SRA data would be named MLP G+S_small_−S.

#### Impact of Data Diversity and Quantity on Model Performance

To analyse the effect of training data diversity on prediction accuracy the following experiments were designed. First, MLP S-S models for sample source prediction were trained with an increasing number of unique SRA studies in the training data, systematically increasing bias diversity. Only SRA studies containing *>* 100 samples for either class were considered. In order to control for training set size, each SRA study was subsampled to 50 samples before training. Six iterations of this training process were conducted starting with one study (i.e. one bias) per class (biopsy vs. cell line). At each step one additional SRA study per class was subsampled ending with six SRA biases and 350 samples in the training set per class. As a control experiment we chose the largest SRA study available for each class to create a training set with a single bias per class. Starting with 50 samples per class in six iterations we subsampled an additional 50 samples ending with 350 samples, thereby assessing the effect on performance that can be attributed to the dataset size. Subsampling and random selection of SRA studies were repeated 10 times with different seeds and each configuration was trained on 10 different seeds, yielding an estimate of uncertainty.

#### Metrics

We report micro and macro accuracy which are equivalent to mean sample accuracy (msa) and mean class accuracy (mca) respectively. Sample accuracy is a measure of absolute performance on the test data. It reports the fraction of correctly classified samples over all classes:

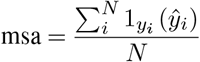

Where *N* is the number of samples, *y* the true label and 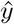 the predicted label, and 1 is the indicator function. Given the large class imbalance in some of our experiments an increase in accuracy in a small class will not be captured by this metric. Average class accuracy, on the other hand, reports the average sample accuracy per class, weighing each class equally and thereby capturing local improvements of the models:

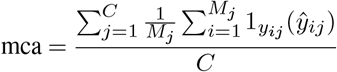

Here, *C* is the number of classes, *M*_*j*_ is the number of samples for class *j*, and *y*_*ij*_ and 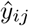 are the true and predicted values, and 1 is the indicator function.

#### Statistical Tests

Accuracy distributions for sex and tissue prediction were tested for statistically significant differences using a t-test (two distributions, scipy.stats.ttest_ind v 1.3.1) or ANOVA (more than two distributions, scipy.stats.f_oneway) with a significance threshold of 0.01.

## RESULTS

### Experimental Setup

This study aims to find the best model for RNA-seq metadata annotation based on gene expression. Three different data sources were selected for which phenotype data was available (Figure 1A). Each of the three data sources comes with a different number of data set biases. Briefly, GTEx is a large homogeneous dataset containing healthy samples following a strict centralized standard protocol. TCGA contains pooled samples from different cancers, disease stages and sequencing centers. Our SRA data is made up of hundreds of individual studies following no centralized standard, containing the largest number of biases of all three data sources. Bias in a test dataset that has not been learned by a model can severely compromise performance. We hypothesized that exposing classification models to a sufficient number of dataset biases will enable them to learn a generalized internal feature representation. Such a model would be able to classify data with previously unseen biases. To test and benchmark our models we selected the classification tasks of (1) tissue of origin of a given RNA-seq sample, (2) biopsy vs. cell-line origin of a sample (i.e. sample source), and (3) sample sex (Figure 1A).

Three different machine learning models were compared (Figure 1B). First, a fully connected ANN (MLP) was tested because of its capability to create novel latent features (see methods for model details). Second, we developed a domain adaptation (DA) approach (Figure 2), a subfield of machine learning dealing with dataset biases. Lastly, the LIN model trained on GTEx data, proposed in Ellis et al. (20), was used as the baseline for all tissue and sex classification experiments.

Models were trained on either GTEx or a mix of GTEx and SRA data and tested on TCGA and SRA data. Uncertainties for MLP and DA models were estimated from 10 training runs with different random seeds (Figure 1B).

### Domain Adaptation Outperforms Other Models on Tissue Classification

We first tested the performance of the LIN, MLP, and DA algorithms to predict the tissue of origin on GTEx (n=5,480), TCGA (n=8,624), and SRA (train n=1,721, test n=1,531) datasets. A subset of 16 tissue labels was chosen that is common to all three data sources (see methods, Supplementary Figure 3, Supplementary Table 3). First, we conducted a single-bias experiment, i.e. MLP G-G (see Nomenclature of Experiments in methods). The nearly perfect score of mean sample accuracy (msa) 0.996 and mean class accuracy (mca) 0.99 (data not shown) confirmed that the MLP yielded highly accurate results when trained and tested on a single-bias dataset (for details on model training, validation, and testing see methods).

### Prediction of SRA Tissue

Metadata prediction on SRA was the most challenging and interesting task due to the potentially large number of different biases in the data source. We re-trained and tested LIN G-S on our datasets and achieved a msa of 0.893 and a mca of 0.765 for the 16 tissues (Figure 3A). Of note is the significantly higher accuracy achieved with LIN G-S compared to the one reported by Ellis et al. (20) (0.519 msa). MLP G-S (msa: 0.872, mca: 0.77) had a higher mca but a lower msa than the corresponding LIN model (Figure 3A). In the next step we investigated the effect of adding bias to the training dataset on prediction performance. In particular, we first predicted SRA tissue from S_small_ data. MLP S_small_-S (msa: 0.894, mca: 0.746) matched the base model’s msa score but performed slightly worse using the mca metric. Similarly, the LIN S_small_-S model matched the msa of LIN G-S but showed an increased performance for mca (msa: 0.893, mca: 0.795) Notably, by only using the small SRA training dataset, we lose the advantage of the large sample size of GTEx. Based on this we hypothesized that by combining SRA and GTEx in the training data, we may be able to leverage both sample size and diversity. The LIN G+S_small_-S model increased its msa to 0.908 and mca to 0.785 which in turn is 1 percentage point (ppt) lower than the LIN S_small_-S model. The two best performing models were MLP G+S_small_-S and DA G+S_small_-S, outperforming LIN G-S on msa by 2.5 ppts and mca 5.5 ppts (MLP G+S_small_-S msa: 0.915, mca: 0.817 and DA G+S_small_-S msa: 0.922, mca: 0.821). No significant d ifference i n the mean performance was detected between these two models (msa p-val*>*0.01, mca p-val*>*0.01, t-test). Crucially, however, DA G+S_small_-S exhibited the lowest standard deviation (std=0.003 for msa and std=0.009 for mca) of all models tested (Supplementary Table 6). For this reason DA G+S_small_-S was considered the best model for the prediction of tissue on the highly heterogeneous SRA test data, increasing the msa score by 1.5% compared to LIN G+S_small_-S and mca by 3.3% compared to LIN S_small_-S, the best performing linear models for the respective metrics.

**Fig. 3.**
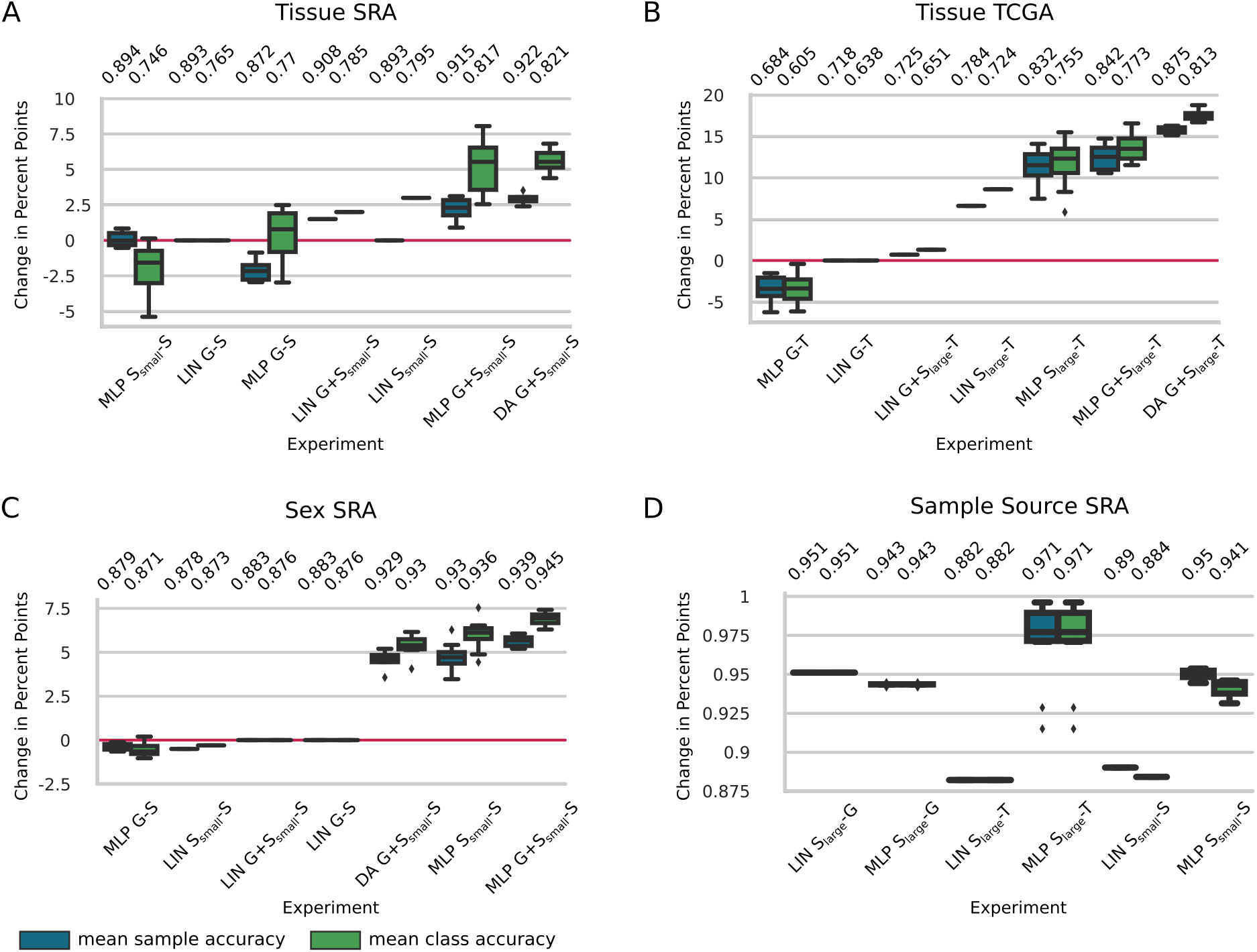
Phenotype Prediction Results for A,B) prediction of tissue of origin on SRA and TCGA (16 classes) C) prediction of sex on SRA (2 classes) and D) sample source (2 classes) on SRA data. Indices ‘small’ and ‘large’ refer to the different size of SRA training data used due to splits of the data set in SRA prediction. Box plots represent model uncertainty of ANN based models, estimated from training with different random seeds (n=10). Mean sample accuracy and mean class accuracy were calculated for each seed. For panel A-C) LIN G-X was chosen as the baseline model. Results for these panels are given in change in percentage points compared to the baseline (red line). Experiments are sorted by increasing mean class accuracy. LIN=linear regression, MLP=multilayer perceptron, DA=domain adaptation, G=GTEx, T=TCGA, S=SRA.

### Prediction of TCGA Tissue

Next, model performance on TCGA data was assessed (Figure 3B). The baseline model LIN G-T achieved msa 0.718 and mca 0.638. Applying the MLP model on the same data resulted in a drop of msa and mca of 2.4 and 3.3 ppts, respectively (MLP G-T msa: 0.684, mca: 0.605). For TCGA tissue prediction we used S_large_ for training, essentially doubling the SRA training data (SRA train + SRA test set: n=3,252). LIN S_large_-T improved accuracy by 6.6 ppts for msa and 8.6 ppts for mca to 0.784 and 0.724 respectively. In comparison, MLP S_large_-T increased model performance by 11.4 ppts to 0.832 (by 11.7 ppts to 0.755) for msa (mca) with respect to LIN G-T. Combining GTEx and SRA training data reduced LIN G+S_large_ performance to msa 0.725 and mca 0.651. The best accuracy was achieved by our MLP G+S_large_ (msa: 0.842, mca: 0.773) and DA G+S_large_ (msa: 0.875, mca: 0.813) models. The DA model had thus a 11.6% performance increase for msa and 12.3% increase for mca compared to LIN S_large_-T, the best linear model. In addition to being the top performer, DA G+S_large_-T also was the most robust model for this task, having the lowest variation in its results (std=0.004 for msa and std=0.006 for mca) (Supplementary Table 6).

We repeated the prediction for TCGA with the models trained for SRA tissue prediction (previous section), i.e. on S_small_, which allows us to assess the influence of the amount of bias injection on model performance. Whereas the addition of more SRA data to the training data set had little influence on LIN models (except for a slight increase of ~0.2 ppts for G-S_large_-T), both MLP and DA model accuracies improved significantly (by between 5 and 9 ppts) upon addition of additional SRA data (Supplementary Table 6).

Notably, adding 5,480 GTEx training samples to MLP S_small_ (MLP-S_small_ → MLP G+S_small_) increased msa from 0.748 to 0.764 and msa from 0.688 to 0.716 on the TCGA test set. On the other hand, adding 1,531 SRA samples (MLP-S_small_ → MLP S_large_) increased msa to 0.832 and msa to 0.755, underlining our model’s ability to incorporate multiple biases for better generalization (Supplementary Table 6).

### Multi-Bias Data Enhances Tissue Classification on TCGA

For tissue classification on TCGA, mean class accuracy increased by 16.8 ppts between MLP G-T and MLP G+S_large_-T. This confirms our hypothesis that the homogeneity of the GTEx data did not allow the MLP G-T model to generalize to TCGA data, while the addition of SRA training data in MLP G+S_large_-T resulted in a model with significantly improved generalization. To further investigate this result we took a closer look at the per class accuracy for the TCGA tissue prediction (Supplementary Figure 6). MLP G-T was unable to predict samples for three tissues, namely bone marrow (msa: 0.08), ovary (msa: 0.02) and uterus (msa: 0.07), whereas all our other models achieved accuracies between 0.7 and 1.0 on these tissues. Adding SRA data to the training set enabled the model to achieve per tissue sample accuracy of 1.00, 0.704 and 0.67 for bone marrow, ovary and uterus, respectively. We used principal component analysis (PCA) to visualize the dataset bias for these tissues. Interestingly, the GTEx-ovary and TCGA-ovary data points show little overlap in the PCA plot, while the SRA-ovary data overlaps with GTEx-as well as TCGA-ovary data, forming a ‘bridge’ (Supplementary Figure 7A). Other tissues such as liver (MLP G-T msa: 0.98), on the other hand, show an overlap between the GTEx and TCGA data which is reflected in the consistent accuracy across all models (Supplementary Figures 6 and 7B).

### Improved Sex Prediction with ANNs

For sex classification only genes on the X and Y chromosome were used as input features (d=190). We first tested the trivial case MLP G-G by splitting GTEx into training and test sets, achieving sample and class accuracy of 0.995 (data not shown).

### Prediction of TCGA Sex

Sex phenotype prediction on TCGA data was the only task where we were not able to perform significantly better than the linear model. The baseline LIN G-T as well as the other linear models LIN S_large_-T and LIN G+S_large_-T achieved almost perfect accuracy on the TCGA data (msa/mca 0.989 for LIN G-T and LIN G+S_large_-T, msa 0.988 and mca 0.987 for LIN S_large_-T). Our best model, based on the data annotation provided by MetaSRA, was MLP G+S_large_-T with msa 0.947 and mca 0.945 (Supplementary Figure 8).

### Prediction of SRA Sex

All linear models for the prediction of sex for SRA data achieved an accuracy (msa: 0.883 and mca: 0.876 for LIN G-S and LIN G+S_small_-S, msa: 0.878 and mca: 0.873 for LIN S_small_-S) similar to what was previously reported (msa: 0.863 (20)). The MLP G-S model (msa: 0.879 and mca: 0.871) did, on average, perform worse than all the linear models. While adding SRA data to the training set did not improve the LIN model, it increased the performance of MLP and DA models. DA G+S_small_-S (msa: 0.929 and mca: 0.93), MLP S_small_-S (msa: 0.93 and mca: 0.936) and MLP G+S_small_-S (msa: 0.939 and mca: 0.945) differ statistically (p-val<8e-5, ANOVA). A t-test corroborated that MLP G+S_small_-S is statistically the best model (p-val=0.0066, t-test) with a performance increase of 6.3% for msa and 7.9% for mca compared to the best linear model LIN G-S. Results are shown in (Figure 3C).

According to MetaSRA all our training and testing data for sex prediction on SRA stem from patient biopsies. However, at least two of the largest misclassified SRA studies in the test set are clearly cultured cell lines. For example, SRP056612 is a study on the effect of the coronavirus on cultured kidney and lung cells (39) and SRP045611 is a study involving HEK cells, which lack the Y chromosome but are annotated as male by MetaSRA (40). These are two examples of mislabeled SRA data. Clearly, mislabeled data can compromise classifier accuracy, either by providing the wrong ground truth for training or by reporting the false label at the point of prediction.

### Expression Based Prediction of Sample Source

SRA data stems from multiple different sources, from which we selected the two largest, namely either biopsy or (immortalized) cell lines, whereas GTEx and TCGA data are exclusively from biopsies. Starting from the hypothesis that fundamental differences do show on an expression level, we set out to train LIN and MLP models on SRA data to predict the sample source of SRA, GTEx and TCGA. Of note, while we were able to approximately reproduce the original results for LIN S_small_-G and LIN S_small_-S we were not able to do so for LIN S_small_-T (msa: 0.998 reported in publication (20)). LIN S_large_-G (msa/mca 0.951) did slightly better than MLP S_large_-G (msa and mca of 0.943). MLP S_large_-T achieved msa and mca 0.971, outperforming LIN S_large_-T with (msa and mca of 0.882). MLP S_small_-S achieved msa 0.95 and mca 0.941, outperforming LIN S_small_-S with msa 0.89 and mca of 0.884 (Figure 3D).

### Training Data Diversity Outweighs Quantity

Our experiments on phenotype classification seem to indicate that increased training data diversity might enhance classification performance. To learn more about the relationship between the amount of training data and model performance MLP G-S was trained on an increasingly large subset of the GTEx training data for tissue classification. We observed a limited effect on model performance with increased training dataset size. The msa reaches its peak with one third of the available training data, while the mca saturates at about half of the available training data (Supplementary Figure 9).

To test the effect of bias in the training data, an MLP S_small_-S for sample source classification was trained on an increasing number of biases in the training set. As a control experiment an MLP was trained with the same amount of data but drawn from a single-bias source. We observed a positive correlation between msa and the number of biases in the training set (Figure 4A). Contrary to that, increasing the number of training samples by the same amount but from a single-bias source did not lead to better model performance (Figure 4B), validating our assumptions. Both experiments support our assumption that ANN based models can integrate different biases in the training set and translate them into better model performance compared to other methods.

**Fig. 4.**
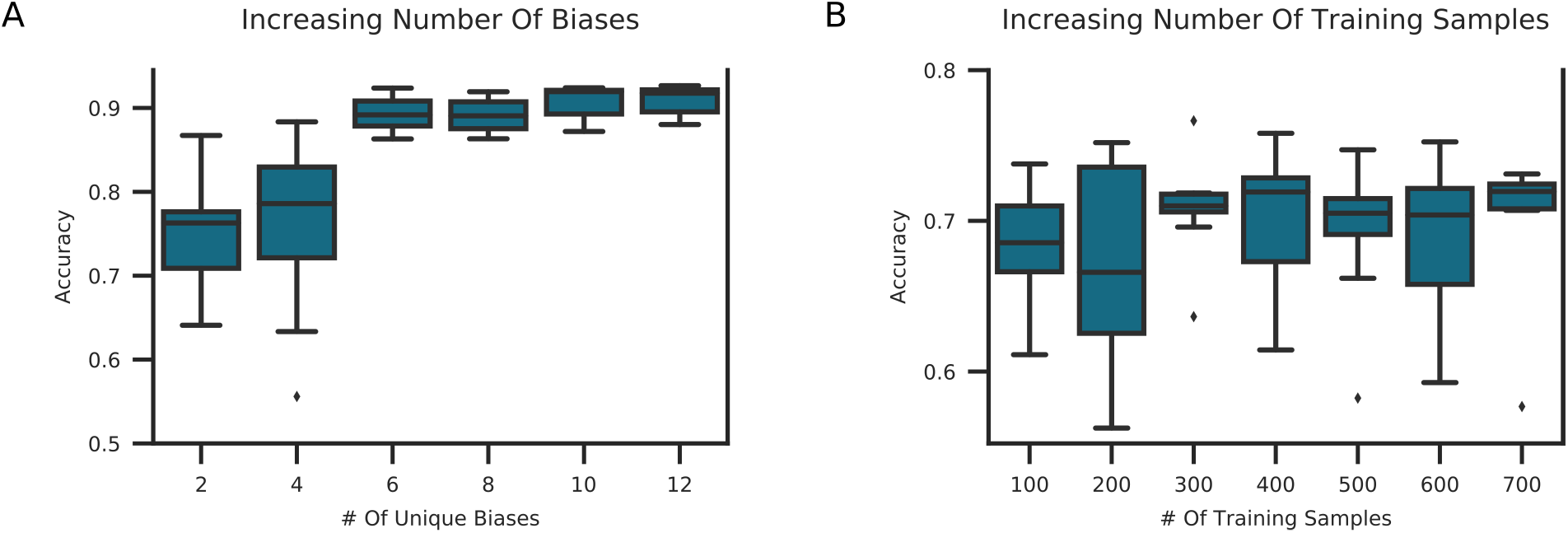
Increasing Bias Vs. Increasing Sample Size in Training Data. A) A MLP S_small_ for sample source prediction on SRA data was trained by randomly sampling an increasing number of SRA studies per class. Each study was subsampled to 50 samples. Studies were drawn from all SRA studies with n *>* 100 for either sample source tissue or cell line. B) To differentiate the effect of increased bias vs. increased sample size, the same model was trained by randomly subsampling the largest available SRA study per class. At each step an additional 50 samples were added to the training set per class. Models were run with 10 different seeds and the mean sample accuracy was computed. Box plots are produced by 10 random sampling iterations. We observe a positive correlation between training data diversity and accuracy.

### Prediction and Availability of Novel Metadata

We have used our best models to predict high-quality metadata for published SRA samples lacking information on tissue, sex, or sample source. Prediction of sex is straightforward because our models were trained on all possible biological categories. For tissue and sample source, however, our models were trained on a subset of all potential classes in the unlabeled data. If, for example, we try to label a sample of a tissue type unknown by the model, the model will force one of the learned classes onto that sample. To deal with this in the best possible way for sample source classification we modified the classification task into one vs. all. Specifically, we first trained a new MLP model to identify the sample source biopsy vs. all other sample sources available in the SRA data as defined by MetaSRA. This model (i.e. MLP S_small_-S) achieved msa 0.947 and mca 0.93 on a test set (data not shown) and MLP S_large_ was subsequently used to identify all of our as of yet unannotated SRA samples of source type biopsy. At a probability cutoff of 0.5 we identified 1,072 new SRA samples as originating from a biopsy.

Second, we extended the tissue classification task to 17 classes by adding a ‘catch-all’ class. To this end, we extended the training data to all GTEx (n=9,366) and SRA (n=6,183) data with tissue labels and assigned the placeholder class for every sample that did not belong to the original set of 16 tissues. That way we ensure that the learned model will not force known classes on every tissue type. With this approach, the DA G+S_small_ model achieved msa 0.912 and mca 0.787 (data not shown). Training and test datasets were subsequently combined to train DA G+S_large_ for annotation prediction of unlabeled SRA samples. We predicted the tissue of origin for all SRA samples of source type biopsy for which no entry on MetaSRA was available (n=2,818).

Third, 8,495 SRA biopsy samples with missing sex information were predicted using MLP G+S_large_. Supplementary Figure 10 shows the true positive rate for each phenotype and each class on the test set. We provide this information such that users can make their own decision on probability cutoffs applied to each class. We provide the full list of all classified SRA samples as well as the probability output of the classifier in the Supplementary Material.

Finally, we used the newly annotated data to improve our models (Figure 1C). To determine the additional tissue training set we chose a probability cutoff of 0.9 and removed all brain samples (n=1,057) to avoid further increase in class imbalance, adding a total of 530 new SRA samples to the training set for tissue prediction (i.e. S_new_). While no increase in performance was observed for TCGA classification, MLP S_small_+S_new_-S for tissue prediction increased in performance compared to MLP S_small_-S from 0.894 to 0.911 (msa) and from 0.746 to 0.798 (mca). DA G+S_large_+S_new_-S achieved the best accuracy of all tissue classification models with msa 0.933 and mca 0.854.

To build a new SRA training dataset for sex all newly annotated SRA samples were added (S_new_=8,495). MLP S_small_+S_new_-S for sex classification i mproved o nly slightly upon the previous best model MLP G+S_small_-S with msa 0.945 and mca 0.948 (compared to msa 0.939 and mca 0.945). For sex classification on TCGA, however, MLP S_large_+S_new_-T yielded sample and class accuracy 0.975, 4.1 ppts higher than the MLP S_large_-T model trained on our default SRA training set (Supplementary Figure 8). We thus successfully identified novel training data and used it in a positive feedback to enhance our models validating the high-quality of the new annotations.

## DISCUSSION

We developed a novel deep-learning based domain adaptation approach for automated bias invariant metadata annotation. To the best of our knowledge this is the first time domain adaptation has been applied to this problem. We were able to outperform the current best model (20) on tissue prediction by 3.3% for SRA and 12.3% for TCGA data on mean class accuracy. We can confirm, as was previously reported (17), that ANNs trained on single-bias training data do not perform better than linear models. Given multi-bias training data, however, we showed that MLPs, and especially our DA algorithm, have an advantage over standard machine learning approaches (e.g. linear regression). Our current models help researchers to verify the sex, tissue and sample type of a RNA-seq sample in the presence of bias. This metadata information is currently rarely given for datasets downloaded from the SRA but can be of crucial importance.

The main strength of our method is its ability to incorporate dataset bias from datasets with only a few samples by applying a siamese network-like architecture. The model learns to ignore bias by repeated exposure to (few) samples in (many) different contexts, i.e. as triplets. In addition, it does not rely on feature selection but uses normalized gene count tables and lets the network learn which features carry important information.

Different types of experiments showed the importance of training models on a multi-bias dataset. First, we showed for every phenotype classification that models which had SRA samples included in the training data performed better than models trained only on GTEx data. For tissue classification we further showed that the effect of adding SRA samples to the training data outweighs adding 3.2x as much GTEx data (MLP S_small_ → MLP S_large_ vs. MLP S_small_ → MLP G-S_small_). Second, for SRA tissue classification we showed that there is a limit of accuracy that can be achieved irrespective of the size of the training set. Our experiment showed that peak accuracy is already reached by using 50% of the available data. Lastly, for sample source classification, we directly compared the relationship between the number of biases in the training data, the number of samples and the model performance. We found a positive correlation between the diversity of the training data and the accuracy achieved by that model.

Lastly, we generated novel metadata for SRA samples using our best performing models, adding over 10,000 new meta-data entries for 8,495 SRA samples. We established a positive feedback loop by re-training the existing models for phenotype prediction by adding the newly annotated data to the training set. Expanding the SRA training data in this way worked especially well for TCGA sex classification where an additional 4.1 ppts in accuracy was achieved. The newly generated metadata is now publicly available and can be used for future research. We see this as a first and important step in the general direction of an effort to make publicly available data more accessible and reusable in an automated way.

We observed some limitations to our DA approach. Our experiments showed that the DA model does not perform as well as the MLP for classification tasks with a low number of classes (e.g. sex). At least for the TCGA tissue classification it seems that a minimum of about 8 classes is needed for the DA model to be able to unfold its full potential consistently. Our experiments indicate that the difference between DA and MLP performance will keep increasing, in favor of the DA model, the more classes we add (Supplementary Figure 11). Adding more tissue classes to our model is an important next step. Another limitation is posed by the need for labeled data to train the bias mapper.

Whereas currently the scope of our predictive models has been limited by the availability of data (e.g. intersecting tissue types between datasets, limited size of datasets), the approach is ready to incorporate more data, biases, classes, and more phenotypes, and there is reason to believe that this will confer increased performance of ANN based models, in particular DA models. At the same time, automated annotation ensures that the vast amount of data, currently lying idle in online repositories and institutional data centers, can indeed be leveraged. We believe that this synergy is capable of producing a large and comprehensive body of annotated biological data that will boost knowledge discovery for biomedical research.

## ACKNOWLEDGEMENTS

We would like to thank the members of the Institute for Medical Systems Biology for helpful discussions and comments. In particular we would like to thank Sergio Oller for help with IT problems and support. In addition, we would like the ZMNH IT for their continuous support. We would like to acknowledge TCGA Research Network (https://www.cancer.gov/tcga) for data provision.

---

**Figure S1.**
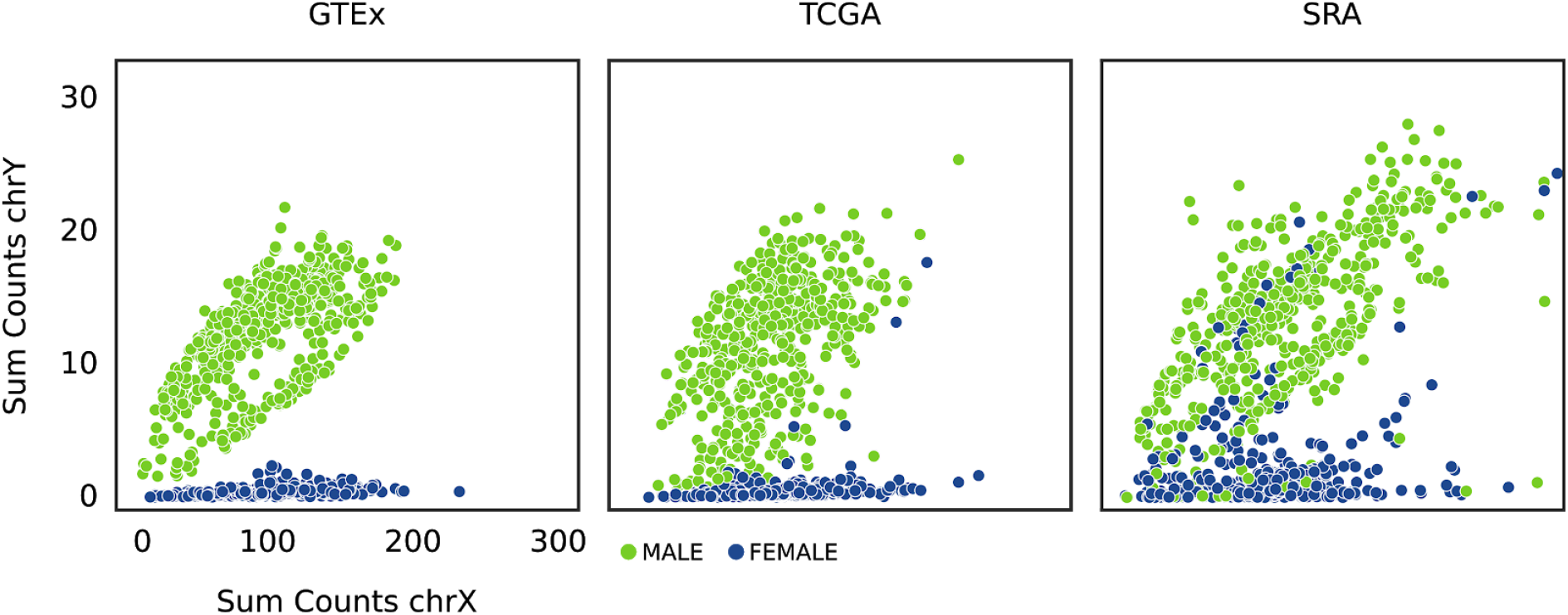
Visualizing Data Set Bias. The three data sources used in this study differ in their homogeneity. Here we drew 1,000 random samples per data source labeled as male (green) or female (blue) and plotted the sum of all gene counts on the Y *vs* the X chromosome. For a highly standardized data source with a solid ground truth we would expect a clear separation of clusters, as we see for GTEx. TCGA shows some overlap in the data due to dataset bias. SRA is the most heterogeneous data source due to multiple data set biases and potentially a less accurate ground truth, as it is a largely uncurated source.

**Figure S2.**
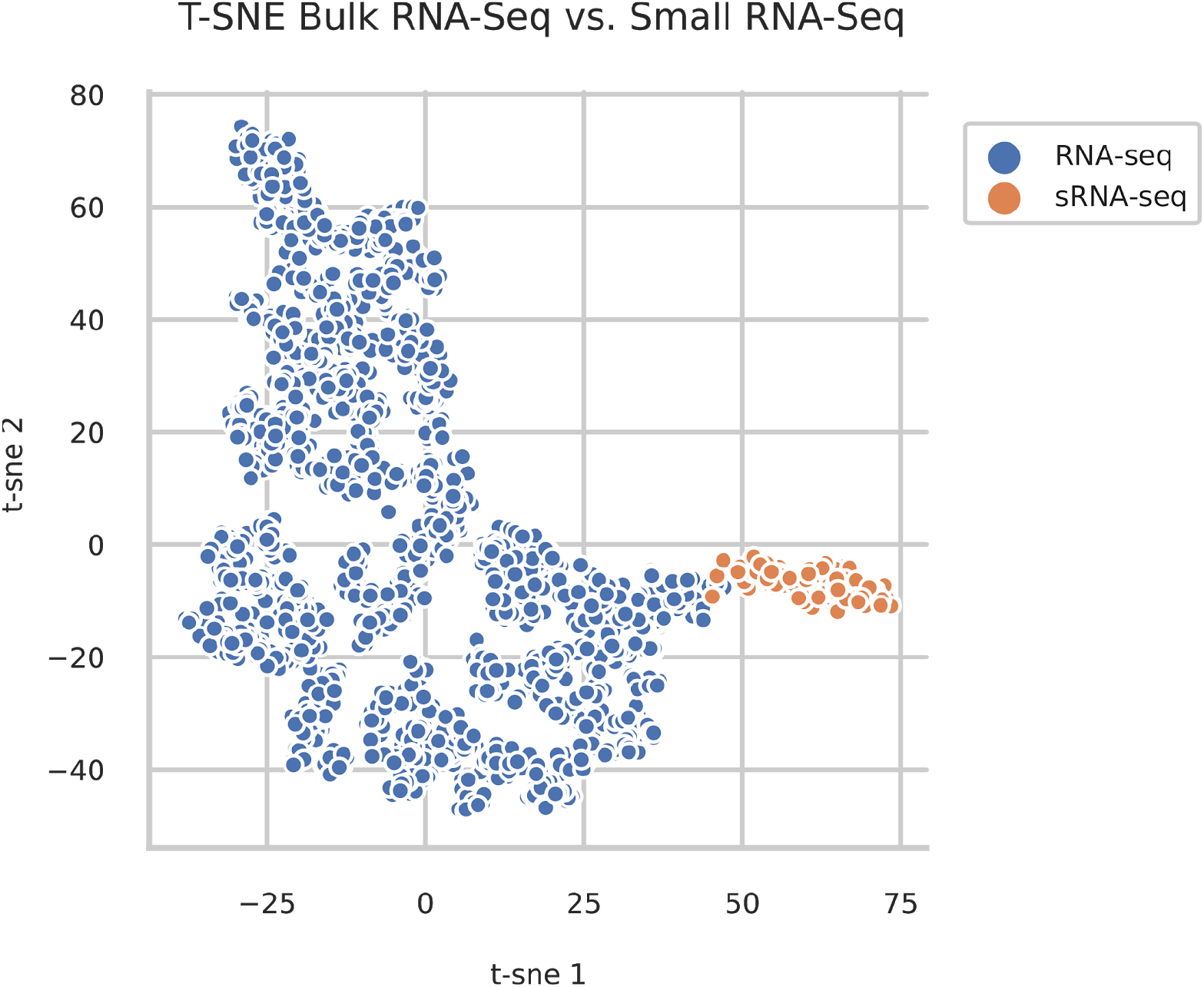
T-SNE on Fraction of Total Gene Count Per Gene Type. The fraction of the total log TPM normalized counts per gene type was calculated for all types that can be associated with mRNA or small RNA. T-SNE was applied on the resulting vectors of fraction per gene type. Samples with their maximum fraction in a gene type belonging to a small RNA category were labeled orange, else blue. The scatter plot shows samples labeled as small RNA-seq all cluster together suggesting a valid approach.

**Figure S3.**
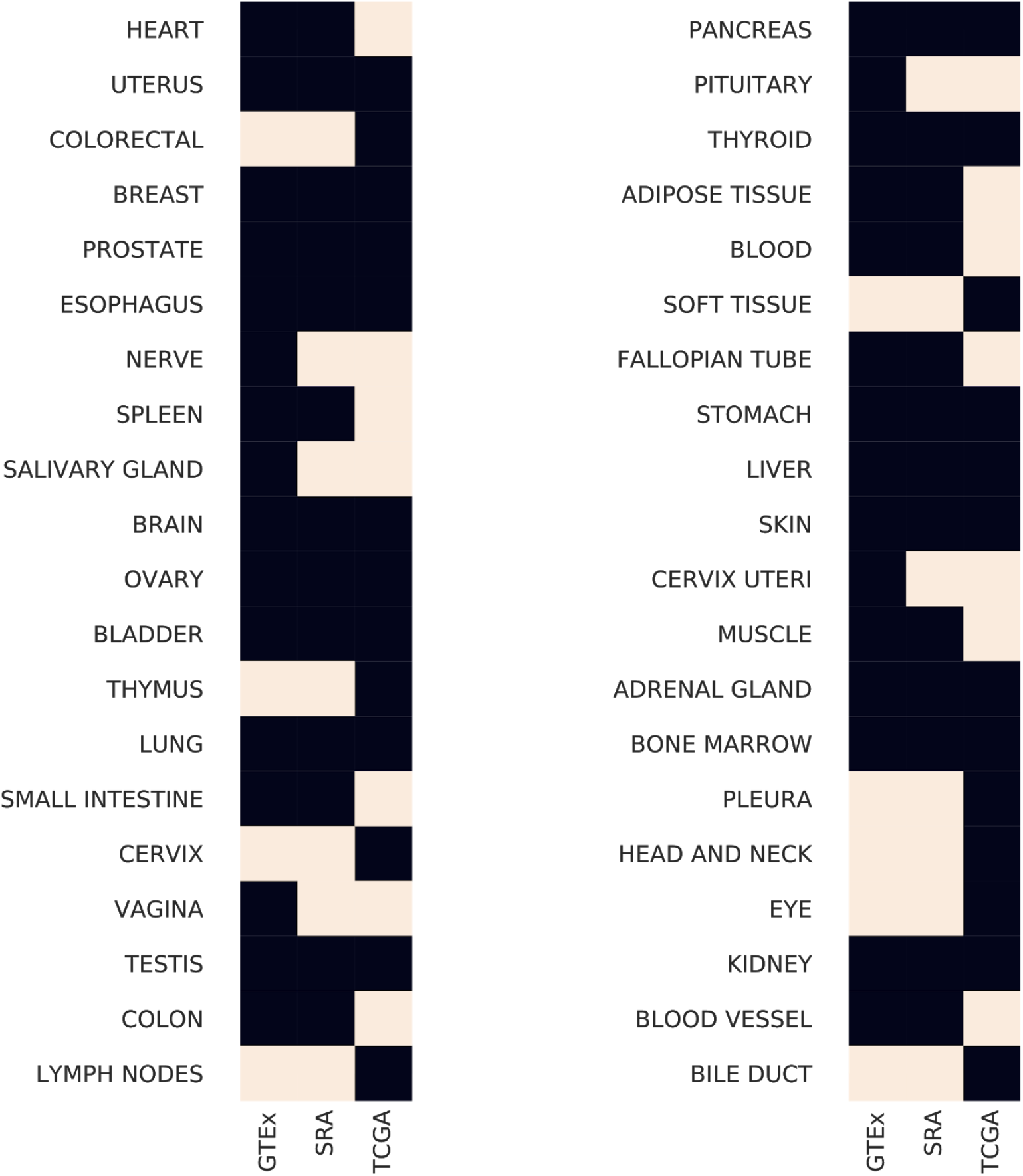
Tissue Label Overlap Between GTEx, TCGA and SRA. GTEx v6 provides samples for 31 tissues and TCGA for 26. MetaSRA provided labels for 26 of the 31 GTEx tissues. This figure depicts the 40 tissues which form the union between the three data sources, a black square indicating that a tissue is present in the respective dataset. 17 Tissues are shared between GTEx, TCGA and SRA, 16 of which were used for tissue prediction.

**Figure S4.**
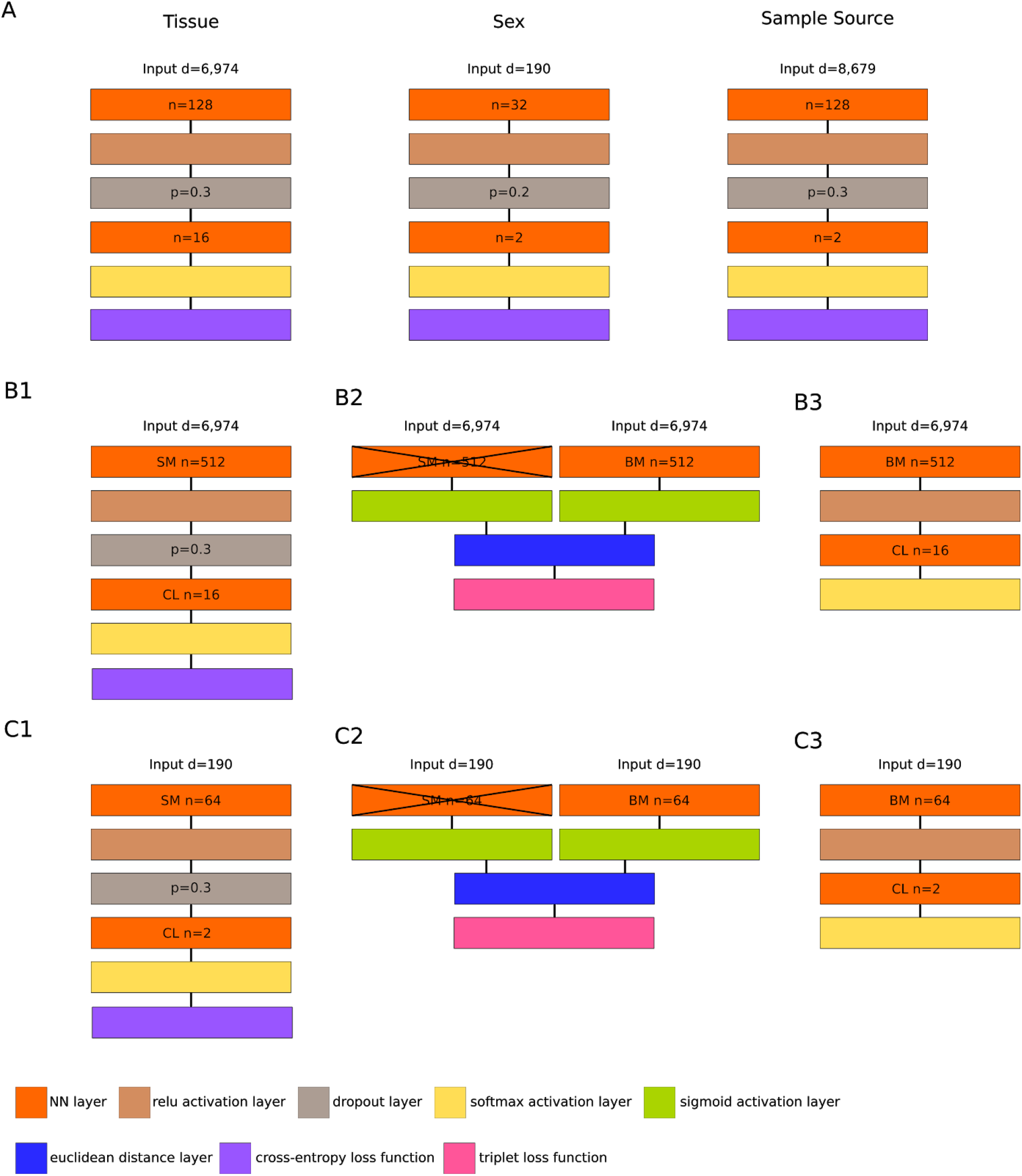
Graphical Representation of Architectures for ANN Based Models. A) MLP models for tissue, sex and sample source, B) are the (1) SM-CL MLP, (2) SM-BM Siamese Network and (3) BM-CL prediction models for tissue and C) sex. Each rectangle represents a layer in the neural network and is colored according to the type of layer that has been used. d=input dimension, n=number of nodes, p=drop out probability, SM=source mapper, BM=bias mapper, CL=classification layer. B2 and C2 show the SM to have frozen weights.

**Figure S5.**
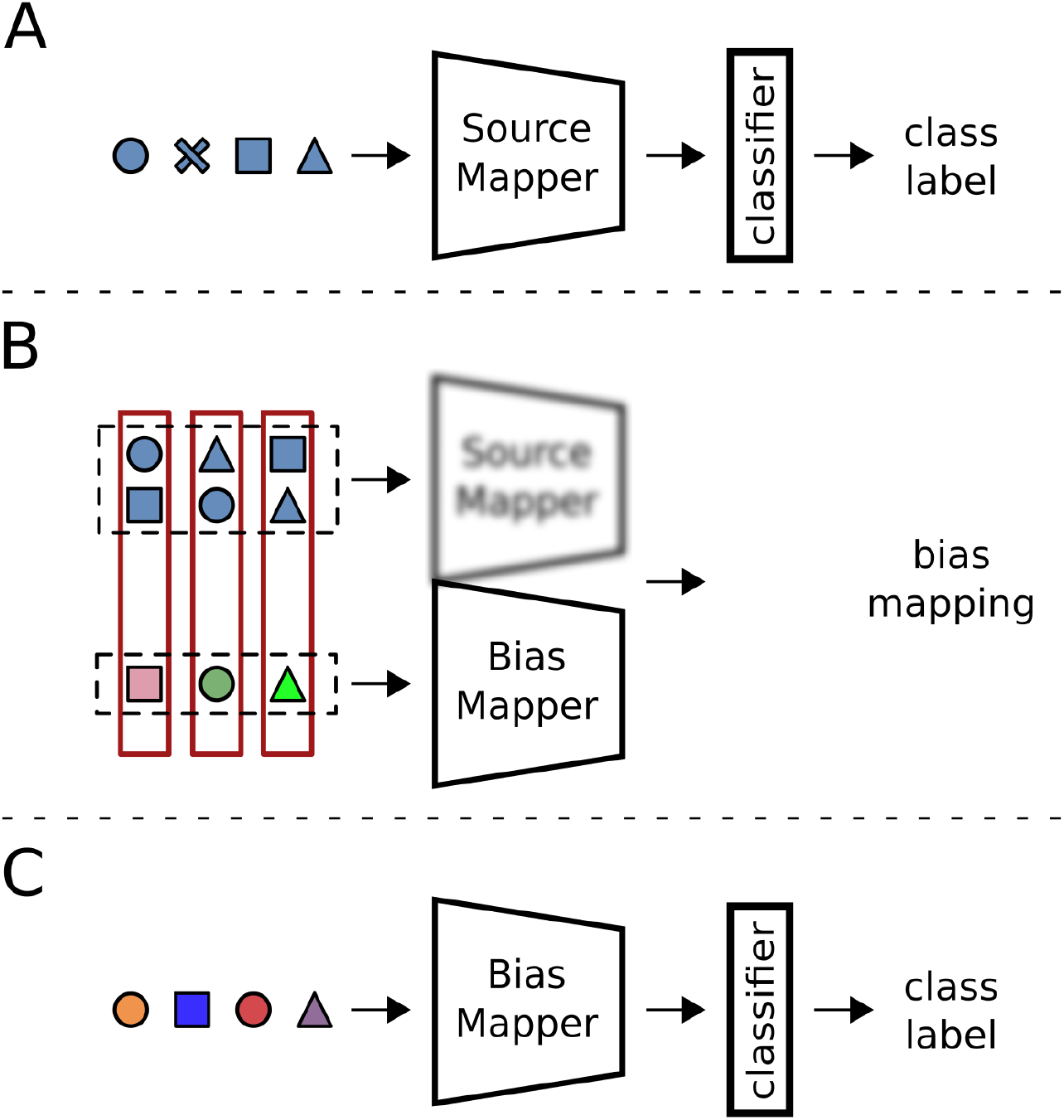
Overview of DA Model. Samples are indicated according to their classes (circles, squares, triangles) and their bias (blue: source domain, other colors: bias domain, target domain). The model is ready for prediction after two training steps: A) A source mapper is trained on single bias data together with a classification layer. B) A bias mapper is created as a duplicate of the source mapper, the weights of the source mapper are fixed. Triplets are passed through the source mapper and bias mapper configuration to learn a bias mapping. C) The bias mapper, equipped with a classification layer, can be used to predict data from previously unseen datasets.

**Figure S6.**
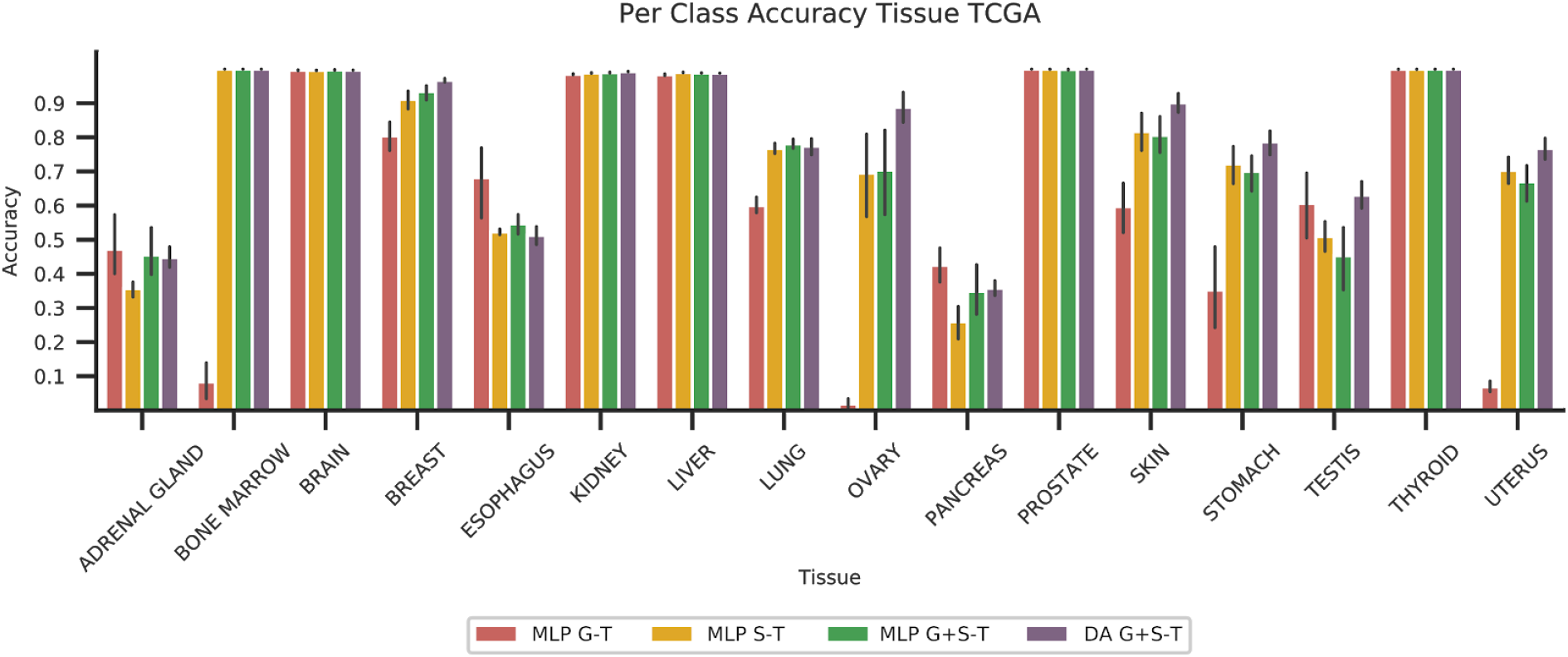
Per Class Accuracy for TCGA Tissue Classification. Mean sample accuracy for each tissue and all ANN based models is shown. The error bar shows the standard deviation across 10 random seeds. The plot demonstrates the varied tissue classification performance of different tissues. For instance, it seems to be difficult to identify adrenal gland or pancreas with any of the models. In particular, the bad classification performance of MLP G-T for bone marrow, ovary and uterus is especially noticeable, along with the observation that performance can be salvaged by addition of (biased) SRA data to the training data set. This highlights the strength of ANN based models in capturing bias from training data.

**Figure S7.**
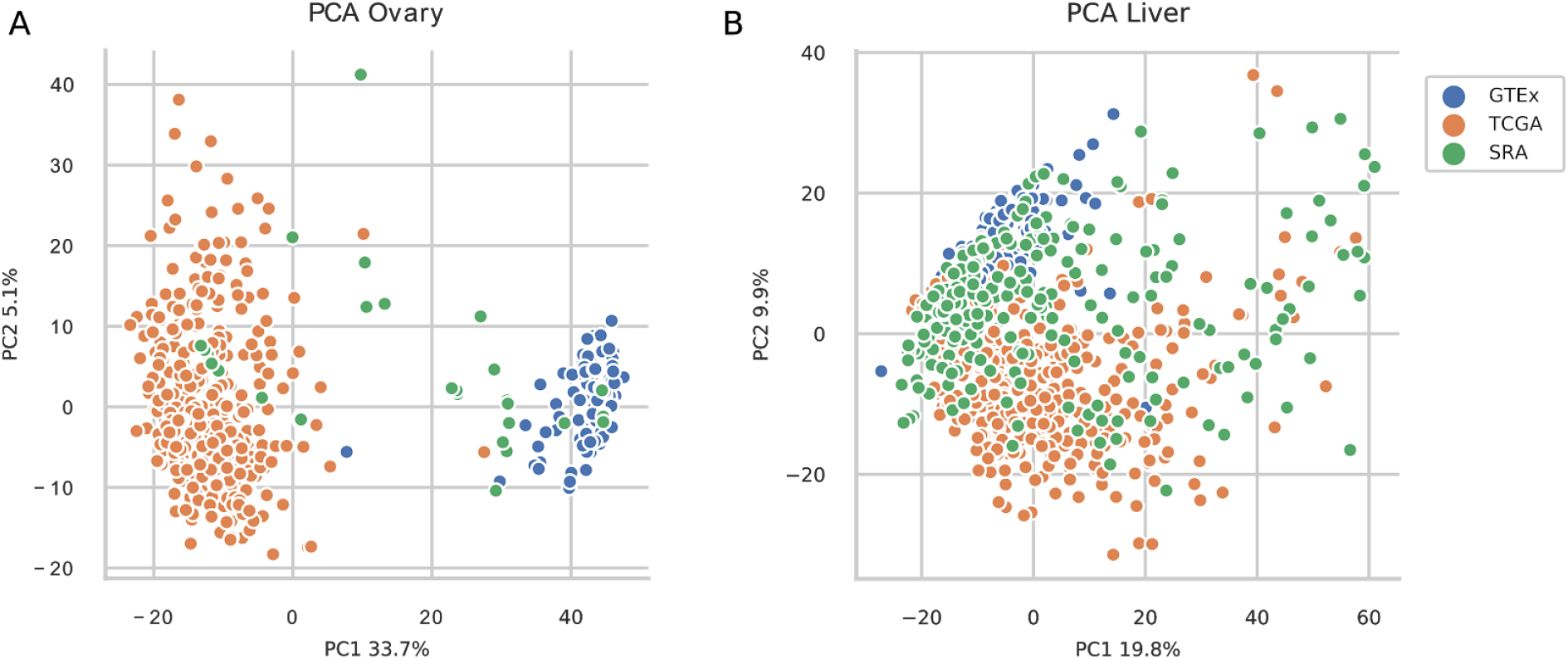
Bias Visualization. Principal Component Analysis on gene expression of available GTEx (blue), TCGA (orange) and SRA (green) samples for (A) ovary and (B) liver tissue. For ovary tissue samples GTEx and TCGA data do not overlap and SRA data is needed for proper model generalization.

**Figure S8.**
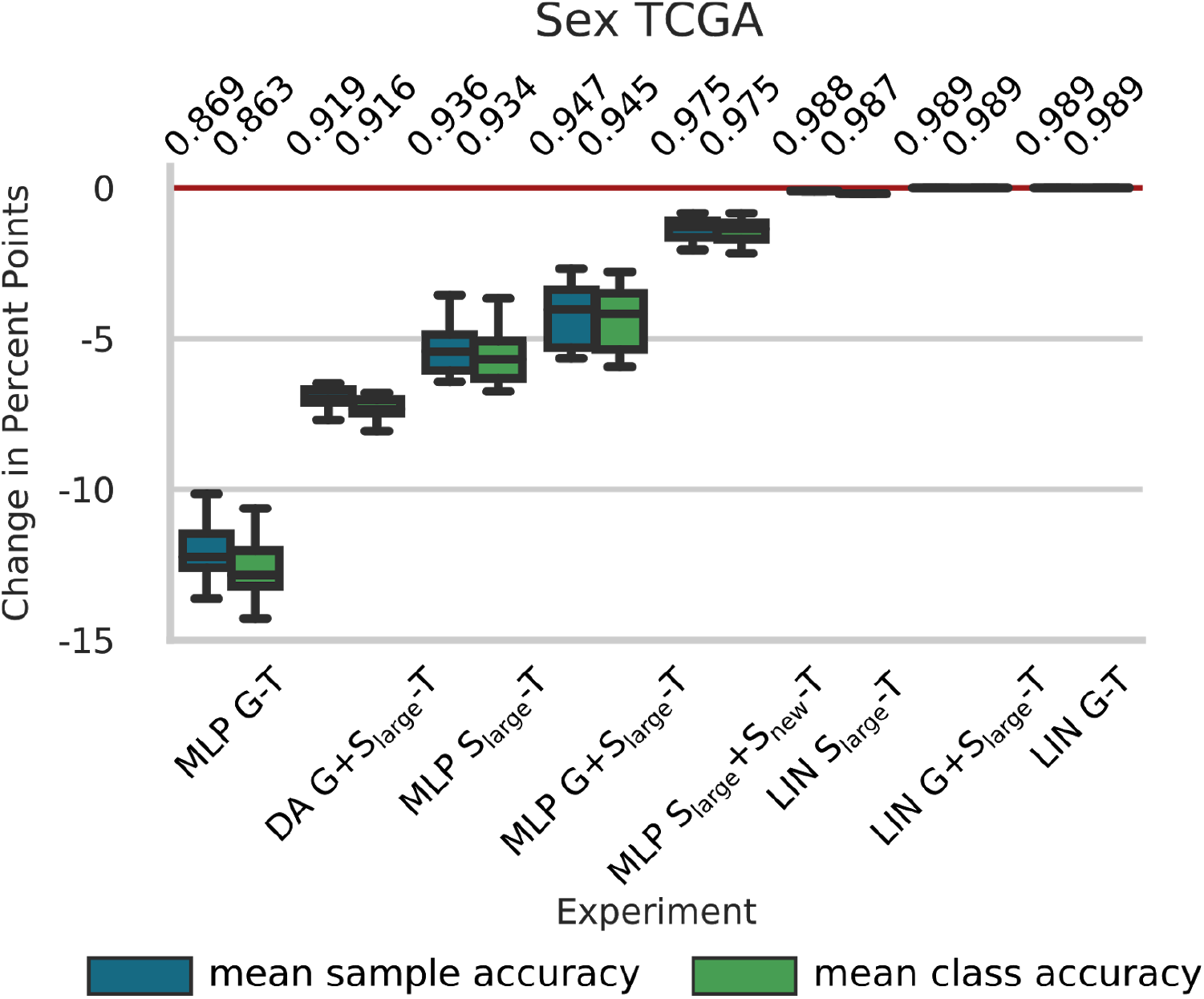
TCGA SEX Results. Results are reported as change in percent points compared with the baseline model LIN G-T. Sample (blue) and class (green) accuracy are shown. LIN=linear model, MLP=multilayer perceptron, DA=domain adaptation, G=GTEx, S=SRA and T=TCGA. ANN based models yielded consistently worse results than the baseline model, until newly annotated data were incorporated into the training set.

**Figure S9.**
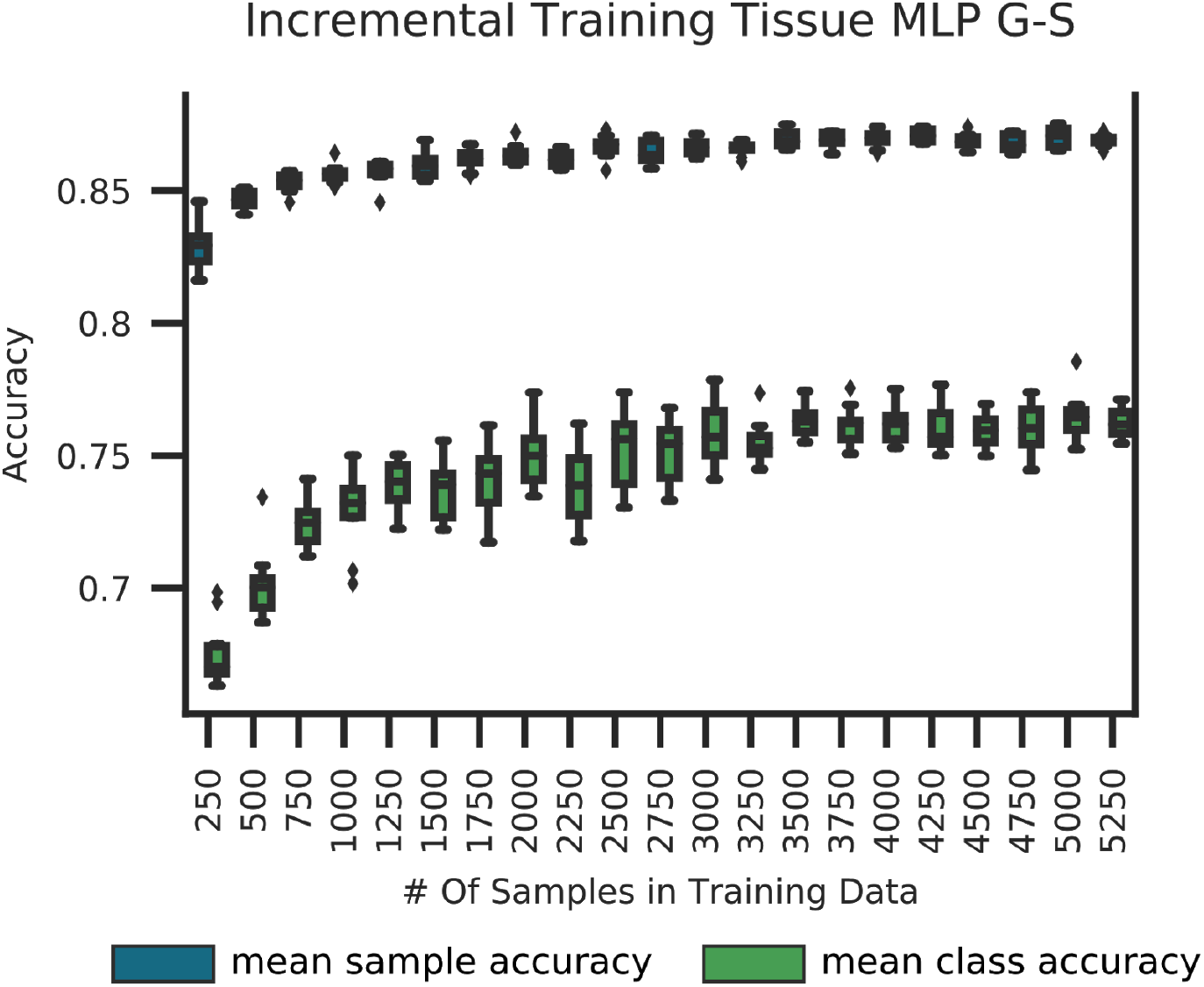
Dependence of Prediction Performance on Increasing Training Data Set Sizes For MLP G-S. MLP models were trained on subsets of the GTEx data for SRA tissue classification on 10 seeds and averaged. At each step the subset was increased by 250 samples. Box Plots from 20 iterations for the msa and mca are shown in blue and green, respectively. Mean sample accuracy reaches its peak with only 25% of the training data while 50% of the data is sufficient for the mean class accuracy to saturate.

**Figure S10.**
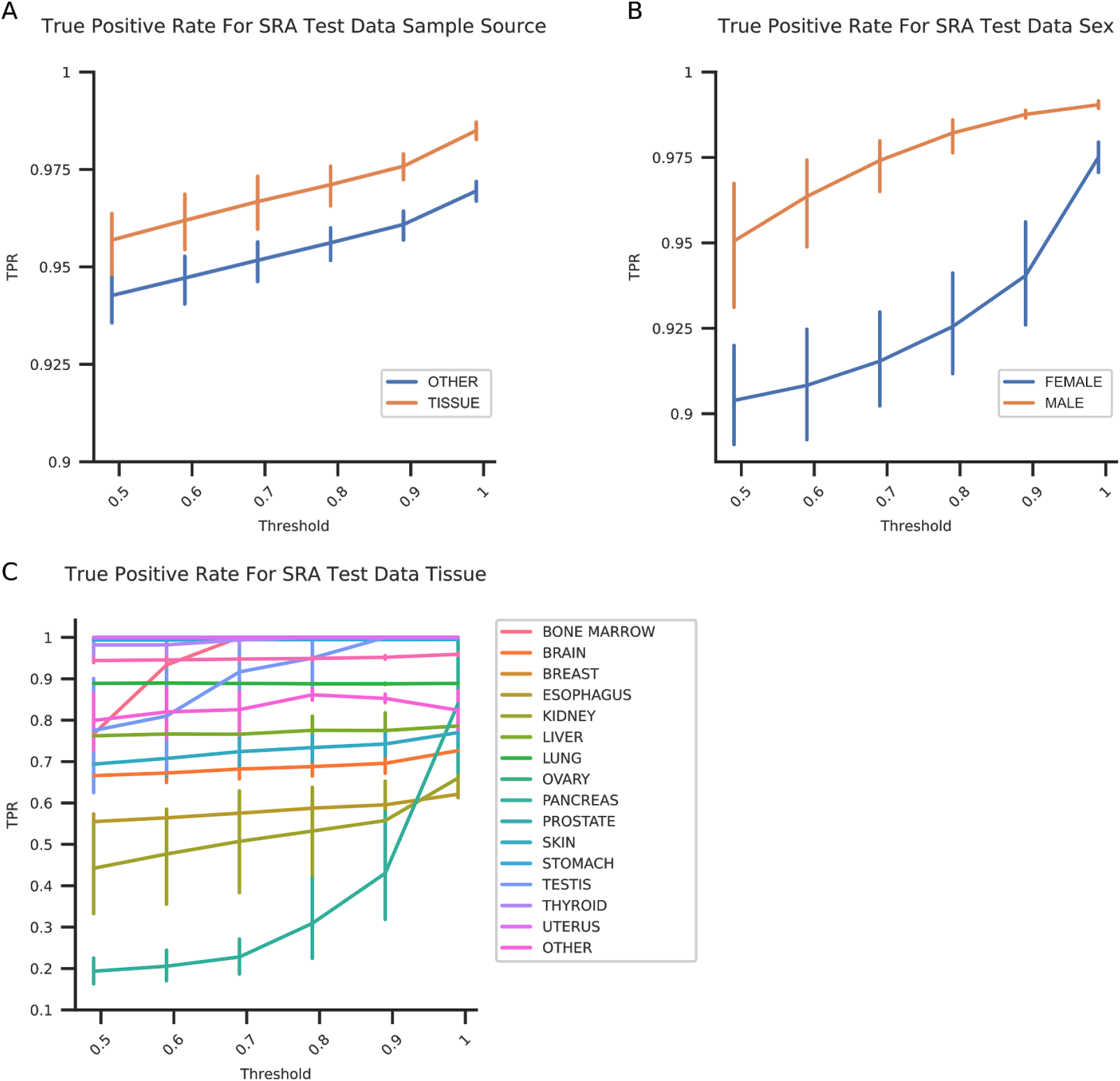
True Positive Rate for Test Data Predicted With Annotation Models. (A) Sample source, (B) sex and (C) tissue classification.

**Figure S11.**
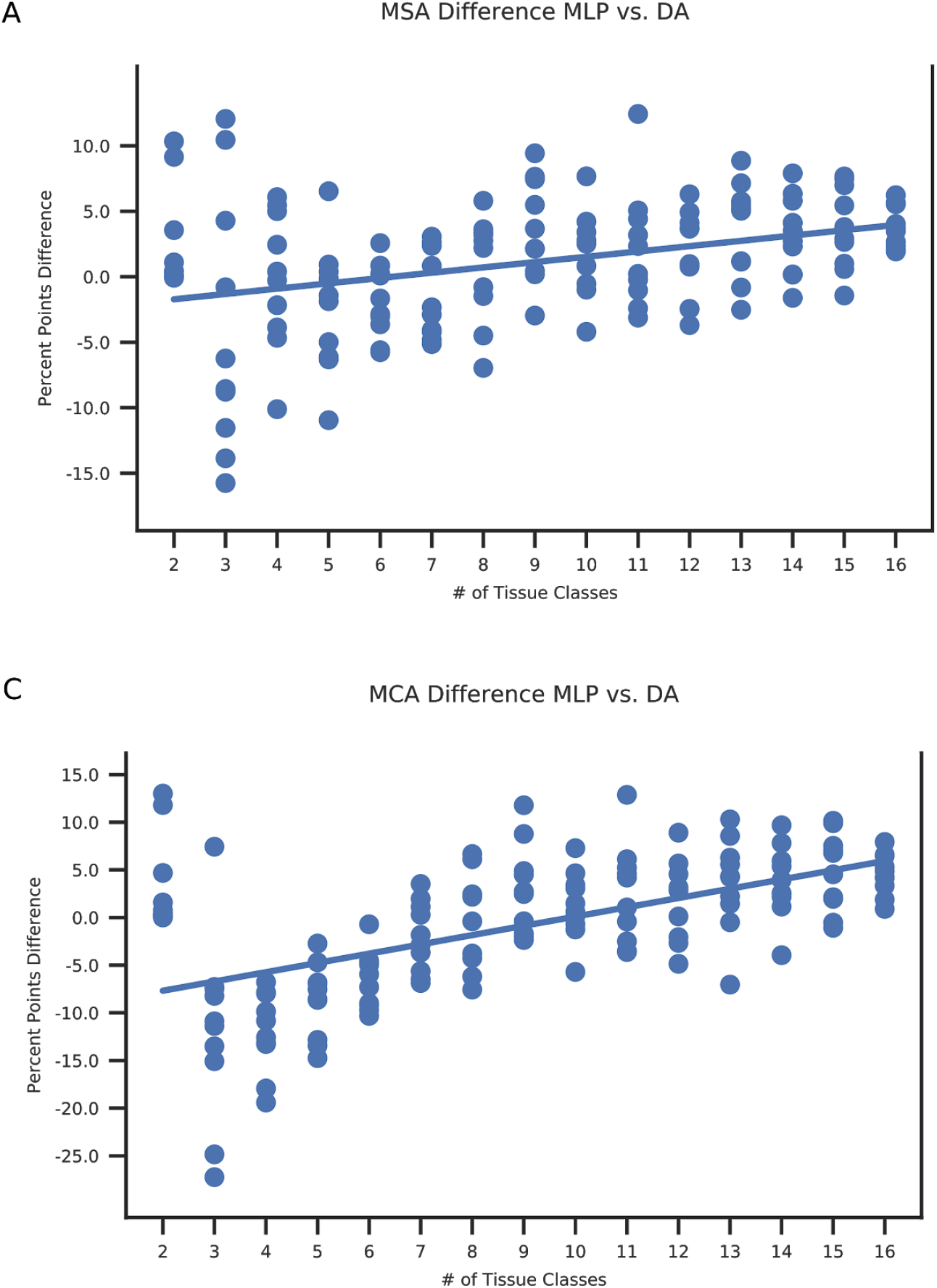
Relationship Between Number of Classes And DA Performance in DA G+S-T. The 16 tissues were sorted by sample size in GTEx, at each step one tissue was added to the classification problem, starting with the largest two. MLP and DA were trained as described above for 10 seeds each and tested on TCGA data. The mean sample accuracy for each seed (top panel) or mean class accuracy (bottom panel) are shown. Each dot shows the difference in accuracy (DA-MLP) at each step for each seed. Seaborn’s regplot was used to fit a regression line. While, on average, MLP performs better for lower number of classes, the performance gain by the DA model with respect to MLP increases with the number of classes.

**Table S1:** Mapping from GTEx tissue names to MetaSRA tissue names.

| GTEx | MetaSRA |
| --- | --- |
| ovary | female gonad |
| skin | anatomical skin |
| thyroid | thyroid gland |
| prostate | prostate gland |
| bladder | urinary bladder |
| cervix uteri | uterine cervix |

**Table S2:**
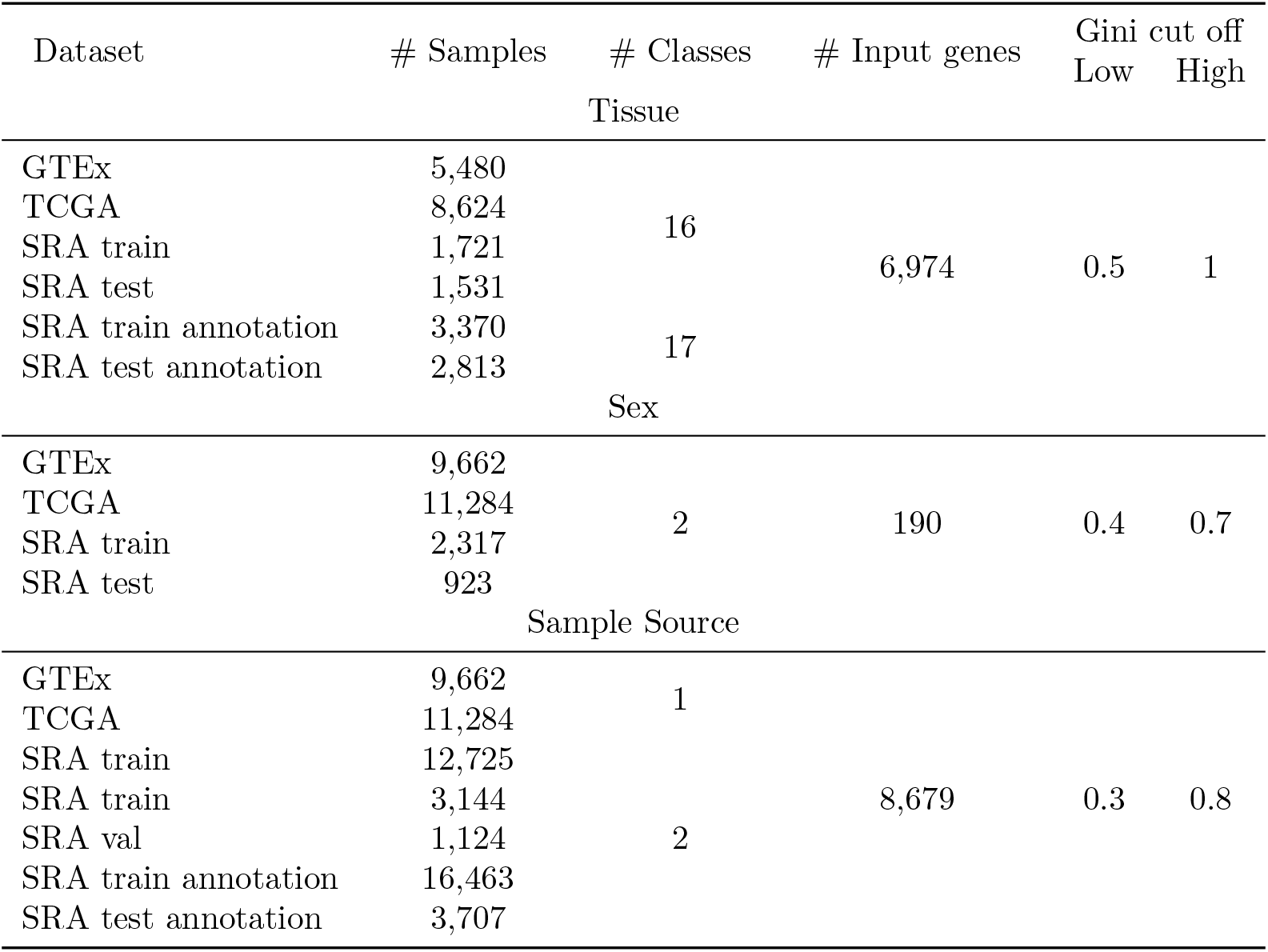
Summary of the datasets used for each phenotype after pre-processing.

**Table S3:** Number of samples per class for phenotype classification experiments.

|  | GTEX | TCGA | SRA train | SRA test |
| --- | --- | --- | --- | --- |
| Tissue |  |  |  |  |
| Adrenal gland | 159 | 266 | 14 | 5 |
| Bone marrow | 102 | 126 | 77 | 90 |
| Brain | 1,409 | 707 | 508 | 770 |
| Breast | 218 | 1246 | 123 | 30 |
| Esophagus | 790 | 198 | 35 | 5 |
| Kidney | 36 | 1030 | 94 | 88 |
| Liver | 136 | 424 | 111 | 134 |
| Lung | 374 | 1156 | 228 | 72 |
| Ovary | 108 | 430 | 23 | 12 |
| Pancreas | 197 | 183 | 17 | 5 |
| Prostate | 119 | 558 | 123 | 49 |
| Skin | 974 | 473 | 238 | 198 |
| Stomach | 204 | 453 | 25 | 11 |
| Testis | 203 | 156 | 14 | 18 |
| Thyroid | 361 | 572 | 51 | 32 |
| Uterus | 90 | 646 | 40 | 12 |
| Sex |  |  |  |  |
| Male | 6,036 | 5,395 | 1,246 | 575 |
| Female | 3,326 | 5,889 | 1,071 | 348 |
| Sample source |  |  |  |  |
| Cell line | 9,662 | 11,284 | 7,108 | 1,950 |
| Biopsy | - | - | 5,617 | 1,194 |

**Table S4:** Hyperparameters considered during model tuning and their initial range.

| Hyperparameter | Range | Sampling mode |
| --- | --- | --- |
| # Layers | [0,3] | linear |
| # Nodes per layer | [32,512] | linear |
| Batch size | [16,32,64] | step |
| Learning rate | [1e-4, 1e-2] | log |
| Optimizer | [Adam, SGD] | binary |
| Drop out | [0.1,0.2,0.3] | step |
| Gini cut off | manually | manually |

**Table S5:** Summary of the hyperparameters used for each model.

| Model | # Nodes | Dropout rate | Learning rate | Margin |
| --- | --- | --- | --- | --- |
| MLP Tissue | 128 | 0.3 | 0.0002 | - |
| MLP Sex | 32 | 0.2 | 0.0024 | - |
| MLP Sample Source | 128 | 0.3 | 0.0002 | - |
| DA SM-CL Tissue | 512 / 16 | 0.3 | 0.0001 | - |
| DA SM-BM Tissue | 512 / 512 | - | 0.0005 | 5 |
| DA SM-CL Sex | 64 / 2 | 0.3 | 0.0001 | - |
| DA SM-BM Sex | 64 / 64 | - | 0.0005 | 3 |

For every model 1 hidden layer was used, batch size was 64, trained epochs were 10 and the optimizer used Adam.

**Table S6:**
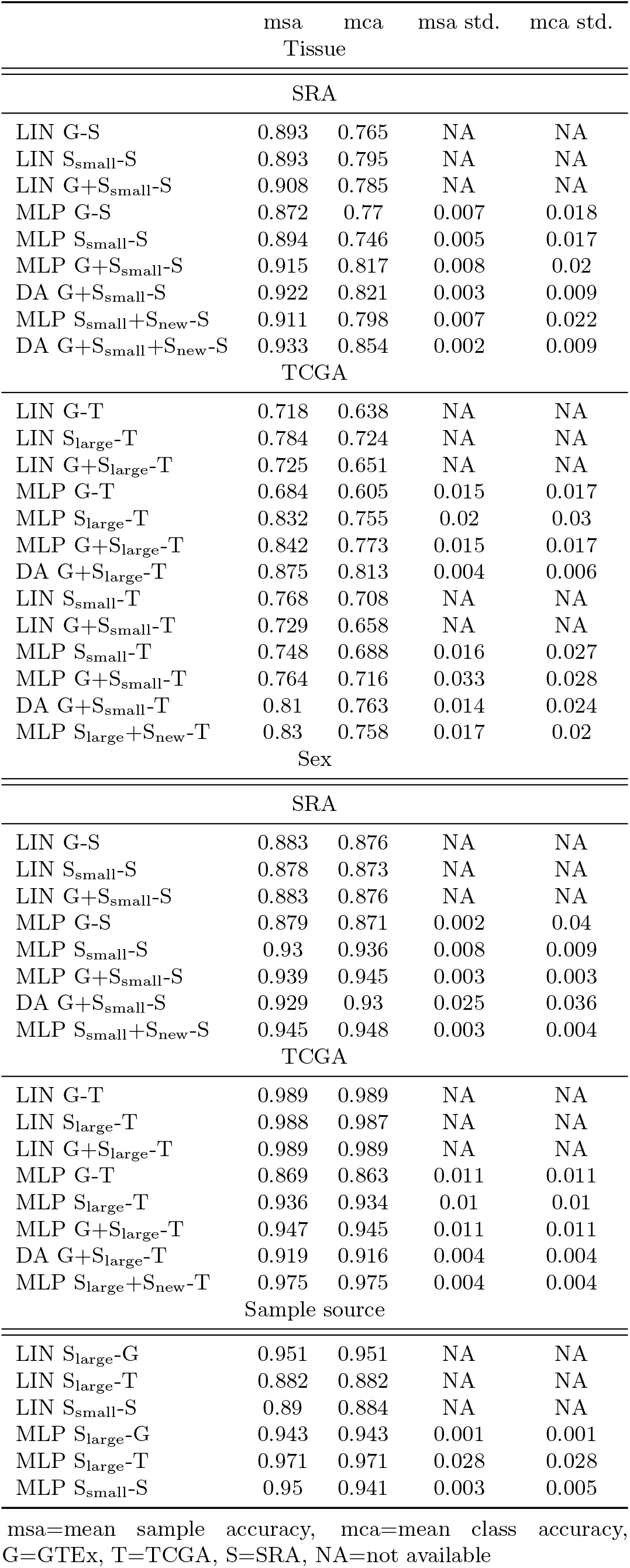
Sample and class accuracy given are the mean over n=10 seeds

|  | msa | mca | msa std. | mca std. |
| --- | --- | --- | --- | --- |
| Tissue |  |  |  |  |
| SRA |  |  |  |  |
| LIN G-S | 0.893 | 0.765 | NA | NA |
| LIN S<sub>small</sub>-S | 0.893 | 0.795 | NA | NA |
| LIN G+S<sub>small</sub>-S | 0.908 | 0.785 | NA | NA |
| MLP G-S | 0.872 | 0.77 | 0.007 | 0.018 |
| MLP S<sub>small</sub>-S | 0.894 | 0.746 | 0.005 | 0.017 |
| MLP G+S<sub>small</sub>-S | 0.915 | 0.817 | 0.008 | 0.02 |
| DA G+S<sub>small</sub>-S | 0.922 | 0.821 | 0.003 | 0.009 |
| MLP S<sub>small</sub>+S<sub>new</sub>-S | 0.911 | 0.798 | 0.007 | 0.022 |
| DA G+S<sub>small</sub>+S<sub>new</sub>-S | 0.933 | 0.854 | 0.002 | 0.009 |
| TCGA |  |  |  |  |
| LIN G-T | 0.718 | 0.638 | NA | NA |
| LIN S<sub>large</sub>-T | 0.784 | 0.724 | NA | NA |
| LIN G+S<sub>large</sub>-T | 0.725 | 0.651 | NA | NA |
| MLP G-T | 0.684 | 0.605 | 0.015 | 0.017 |
| MLP S<sub>large</sub>-T | 0.832 | 0.755 | 0.02 | 0.03 |
| MLP G+S<sub>large</sub>-T | 0.842 | 0.773 | 0.015 | 0.017 |
| DA G+S<sub>large</sub>-T | 0.875 | 0.813 | 0.004 | 0.006 |
| LIN S<sub>small</sub>-T | 0.768 | 0.708 | NA | NA |
| LIN G+S<sub>small</sub>-T | 0.729 | 0.658 | NA | NA |
| MLP S<sub>small</sub>-T | 0.748 | 0.688 | 0.016 | 0.027 |
| MLP G+S<sub>small</sub>-T | 0.764 | 0.716 | 0.033 | 0.028 |
| DA G+S<sub>small</sub>-T | 0.81 | 0.763 | 0.014 | 0.024 |
| MLP S<sub>large</sub>+S<sub>new</sub>-T | 0.83 | 0.758 | 0.017 | 0.02 |
| Sex |  |  |  |  |
| SRA |  |  |  |  |
| LIN G-S | 0.883 | 0.876 | NA | NA |
| LIN S<sub>small</sub>-S | 0.878 | 0.873 | NA | NA |
| LIN G+S<sub>small</sub>-S | 0.883 | 0.876 | NA | NA |
| MLP G-S | 0.879 | 0.871 | 0.002 | 0.04 |
| MLP S<sub>small</sub>-S | 0.93 | 0.936 | 0.008 | 0.009 |
| MLP G+S<sub>small</sub>-S | 0.939 | 0.945 | 0.003 | 0.003 |
| DA G+S<sub>small</sub>-S | 0.929 | 0.93 | 0.025 | 0.036 |
| MLP S<sub>small</sub>+S<sub>new</sub>-S | 0.945 | 0.948 | 0.003 | 0.004 |
| TCGA |  |  |  |  |
| LIN G-T | 0.989 | 0.989 | NA | NA |
| LIN S<sub>large</sub>-T | 0.988 | 0.987 | NA | NA |
| LIN G+S<sub>large</sub>-T | 0.989 | 0.989 | NA | NA |
| MLP G-T | 0.869 | 0.863 | 0.011 | 0.011 |
| MLP S<sub>large</sub>-T | 0.936 | 0.934 | 0.01 | 0.01 |
| MLP G+S<sub>large</sub>-T | 0.947 | 0.945 | 0.011 | 0.011 |
| DA G+S<sub>large</sub>-T | 0.919 | 0.916 | 0.004 | 0.004 |
| MLP S<sub>large</sub>+S<sub>new</sub>-T | 0.975 | 0.975 | 0.004 | 0.004 |
| Sample source |  |  |  |  |
| LIN S<sub>large</sub>-G | 0.951 | 0.951 | NA | NA |
| LIN S<sub>large</sub>-T | 0.882 | 0.882 | NA | NA |
| LIN S<sub>small</sub>-S | 0.89 | 0.884 | NA | NA |
| MLP S<sub>large</sub>-G | 0.943 | 0.943 | 0.001 | 0.001 |
| MLP S<sub>large</sub>-T | 0.971 | 0.971 | 0.028 | 0.028 |
| MLP S<sub>small</sub>-S | 0.95 | 0.941 | 0.003 | 0.005 |

msa=mean sample accuracy, mca=mean class accuracy, G=GTEX, T=TCGA, S=SRA, NA=not available

